# Dev-Atlas: A reference atlas of functional brain networks for typically developing adolescents

**DOI:** 10.1101/2024.08.20.608658

**Authors:** Gaelle E. Doucet, Callum Goldsmith, Katrina Myers, Danielle L. Rice, Grace Ende, Derek J. Pavilka, Marc Joliot, Vince Calhoun, Tony W. Wilson, Lucina Q. Uddin

## Abstract

Adolescence is a critical period for neural changes, including maturation of the brain’s cognitive networks, but also a period of increased vulnerability to psychopathology. It is well accepted that the brain is functionally organized into multiple interacting networks and extensive literature has demonstrated that the spatial and functional organization of these networks shows major age-related changes across the lifespan, but particularly during adolescence. Yet, there is limited option for a reference functional brain atlas derived from typically developing adolescents, which is especially problematic as the reliable and reproducible identification of functional brain networks crucially depends on the use of such reference functional atlases. In this context, we utilized resting-state functional MRI data from a total of 1,391 typically developing youth between the ages of 8 and 17 years to create a new adolescent-specific reference atlas of functional brain networks. We further investigated the impact of age and sex on these networks. Using a multiscale individual component clustering algorithm (MICCA), we identified 24 reliable functional brain networks, classified within six domains: Default-Mode (5 networks), Control (4 networks), Salience (3 networks), Attention (4 networks), Somatomotor (5 networks), and Visual (3 networks). We identified reliable and large effects of age on the spatial topography of these majority of networks, as well as on the functional network connectivity (FNC) between networks. The DMN showed reduced FNC with the other networks with older age. Sex effects were not as widespread. No significant sex-by-age interactions were detected. Overall, we created a novel brain atlas, named Dev-Atlas, focused on a typically developing sample, with the hope that this atlas can be used in future independent developmental network neuroscience studies. Dev-Atlas is freely available to the research community.

## Introduction

Adolescence is a critical period for neural changes, including maturation of the brain’s cognitive networks, but also a period of increased vulnerability to psychopathology (Paus et al., 2008). Structural and functional neuroimaging studies have been instrumental in mapping and improving our understanding of such brain organizational changes throughout development. However, one of the most challenging topics in recent neuroscience has been to quantify inter-subject variability and link it to meaningful neurodevelopmental and/or psychological phenotypes (Marek et al., 2022). One common approach has been to apply a normative brain atlas-based template in order to map major functional brain networks and quantify how their spatial maps and/or functional connectivity (FC) differ among individuals (Gratton et al., 2020). From that point, the association with age, sex and other behavioral variables, or differences between typically developing and clinical participants have been widely investigated (Di Martino et al., 2014; Zhang et al., 2021).

The use of brain atlases has allowed neuroimaging researchers to augment the generalizability and comparability of functional studies across diverse cohorts. Several brain functional atlases are currently available, most of which are based on healthy young adult individuals (Craddock et al., 2012; Doucet et al., 2019; Gordon et al., 2016; Shen et al., 2013; Wig et al., 2014; Yeo et al., 2011). To our knowledge, only very recently (in 2024), developmental-based atlases of brain functional networks have been created (Fu et al., 2024; Hermosillo et al., 2024); however, their validation and use by independent research groups remain to be done. The hope is that these age-specific atlases will help map age-appropriate brain functional parcellation and networks, and reduce inter-subject variability. Indeed, it is well known that the brain functional parcellation is typically more segregated in youth compared with adults, as demonstrated by a larger number of functional brain networks (Fu et al., 2024; Gu et al., 2015). Investigations using ABCD data are beginning to provide longitudinal evidence that youth show greater segregation between networks as they age (Chang et al., 2023). It is therefore possible that the lack of network specificity leads to increased inter-subject variability, or inaccurate brain mapping leading to incorrect interpretation of developmental effects.

Very recently, Fu et al. (2024) created a developmental brain functional atlas (Neuromark_fMRI_3.0) based on resting-state functional MRI (rsfMRI) data collected from 652 young individuals aged 5-21 using the Human Connectome Project-Development sample (HCP-D) (Harms et al., 2018; Somerville et al., 2018). Using a group-based independent component analysis (gICA) with high dimensionality, they were able to identify 67 reproducible independent components (ICs), classified into nine functional systems (subcortical, hippocampal, auditory, sensorimotor, visual, cognitive-control, parietal, default-mode, and cerebellar). They further showed that, while most of these ICs were reproducible and could be detected in other age groups (i.e., infants and older adults), the younger template tended to be more segregated while the older template tended to be more aggregated, and some ICs were specifically identified in only one age group. Overall, this study confirms the need for more validated age-specific brain functional atlases in order to better capture functional features across the lifespan, and particularly during typical development.

In this context, the aim of this study was to construct a new reproducible functional brain atlas focused on typically-developing youth between the ages of 8 and 17 years. For this, we utilized rsfMRI datasets from typically-developing children and adolescent participants from three large developmental projects: the Philadelphia Neurodevelopmental Cohort (PNC) (Satterthwaite et al., 2016; Satterthwaite et al., 2014), the Pediatric Imaging, Neurocognition, and Genetics (PING) study (Jernigan et al., 2016), and the Lifespan Human Connectome Project – Development (HCP-D) (Harms et al., 2018). A total of 1391 individuals were included and 24 reproducible brain functional networks were identified as part of a new functional brain atlas named “Dev-Atlas”. We further tested the impact of age and sex on the spatial maps and functional network connectivity (FNC, i.e., connectivity among overlapping whole brain networks) of each of the derived networks. This atlas is openly accessible for download at https://github.com/BRAICLab/DevAtlas.

## Material and Methods

### Main Cohorts

We utilized datasets from typically-developing adolescent participants from three large developmental projects (**Supplementary Figure S1, Table 1**): the Philadelphia Neurodevelopmental Cohort (PNC) (Satterthwaite et al., 2016; Satterthwaite et al., 2014), the Pediatric Imaging, Neurocognition, and Genetics (PING) study (https://chd.ucsd.edu/research/ping.html) (Jernigan et al., 2016), and the Lifespan Human Connectome Project – Development (HCP-D) (Harms et al., 2018). In each cohort, we selected individuals aged 8 to 17 years for whom both rsfMRI and structural MRI data were available. This selection resulted in a sample of 989 individuals for PNC, 476 for HCP-D, and 229 for PING. Following quality control of the imaging data, 333 participants across cohorts were removed. The total analysis sample comprised 1,391 individuals (**Supplementary Figures S1 & S2**, **Table 1**).

**Table 1:**
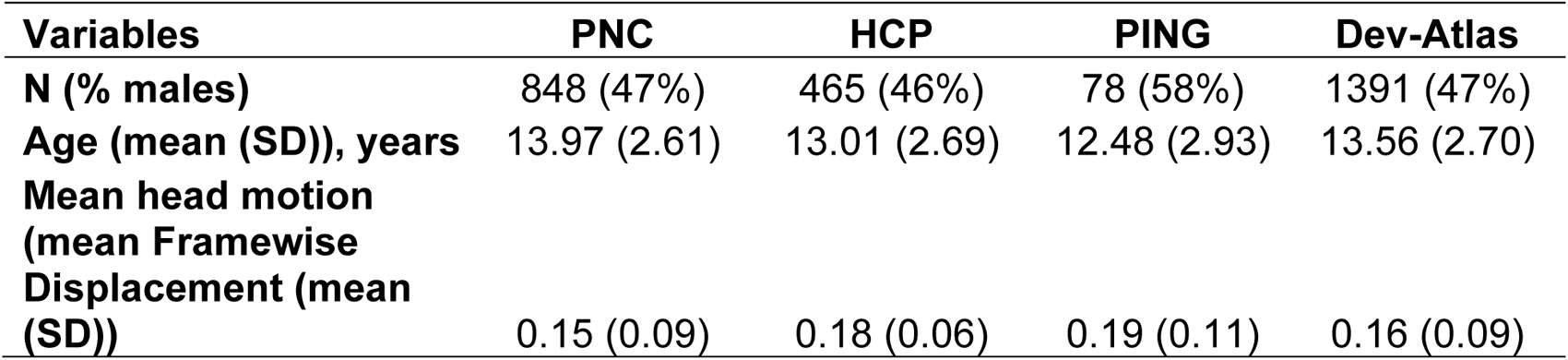
Demographic information of the sample used to create Dev-Atlas.

### BTNRH Cohort

We also used an independent smaller sample of typically developing children and adolescents aged between 8 and 17 years old, collected at Boys Town National Research Hospital (BTNRH), Omaha, NE, USA, for replication (n=214, 53% males, mean age=12.23 (2.63) years). All participants reported no history of psychiatric or neurological disorders, no learning disabilities, no history of head trauma or concussion, no developmental delays, no history or current use of substances, and no metal implants or irremovable ferromagnetic objects that could be an MRI safety concern. The study was approved by the local institutional review board (IRB) and parents consented to have their children participate in the research, while the children provided written assent.

### Resting-State fMRI Acquisition and Preprocessing

In the PNC cohort, rsfMRI data were acquired on a 3T Siemens TIM Trio scanner. The main acquisition parameters were: TR/TE=3000/32 msec, flip angle=90◦; FOV=192 mm×192 mm; voxel-size = 3 × 3 × 3 mm^3^, acquisition time=6:18 min, number of volumes: 124. More information is provided in Satterthwaite et al. (2014).

In the HCP-D cohort, rsfMRI data were acquired on a 3T Siemens Trio scanner. The main acquisition parameters were: TR/TE =800/37 msec, 32 axial slices, flip angle = 52◦; FOV= 208 mm×208 mm; voxel-size = 2 × 2 × 2 mm^3^, acquisition time = 6:40 min, number of volumes: 478. When multiple rsfMRI datasets were available for each participant, the first one was selected for analyses. More information on the MRI sequences can be found in Harms et al 2019.

In the PING cohort, rsfMRI data were acquired on a 3T Siemens TIM Trio scanner. Main acquisition parameters were TR/TE=3000/30 msec, flip angle=90◦, the number of volumes varied between 128 and 300. More information on the MRI sequences can be found in Jernigan et al (2016).

In the BTNRH sample, rsfMRI data were acquired on a 3T Siemens Prisma scanner, using the following acquisition parameters: TR/TE=480/29.2 msec, 56 axial slices, flip angle = 44◦; FOV= 248 mm×248 mm; voxel-size = 3 × 3 × 3 mm^3^, number of volumes varied between: 700 and 775.

The rsfMRI data were preprocessed in an identical fashion for all cohorts using *fMRIprep* v.23.0.2, followed by a final step for denoising using the Nilearn toolbox (nilearn.interfaces.fmriprep.load_confounds). The strategy used was *ica_aroma*, and global signal regression was also applied, following the recommendations by Ciric et al. (2017) to minimize the impact of head motion on functional connectivity (Satterthwaite et al., 2019). More details are provided in **Supplementary Material**.

Across the three main cohorts, we excluded 145 individuals for issues related to the preprocessing pipeline (e.g., unreadable DICOM files) and 158 individuals who exceeded at least one of three excessive head movement exclusion criteria: mean framewise displacement (FD) >0.5 mm, max motion >3 mm, and/or ≥50% of volumes with a FD >0.5 mm (**Supplementary Figure S1**).

### Resting-state Network Identification

We used a process validated by Naveau et al. (2012) to identify reliable brain functional networks, which we also previously used to create a brain atlas for older adults (Doucet et al., 2020). We analyzed the main and replication (BTNRH) cohorts, separately. First, for each individual within each cohort, single-subject ICA with random initialization were conducted, using the Multivariate Exploratory Linear Optimized Decomposition into Independent Components (MELODIC) software, version 3.15, included in the FMRIB Software Library (FSL) (Smith et al., 2004). The number of independent components (ICs) was estimated by Laplace approximation (Minka, 2000). A symmetric approach of the FastICA algorithm (Hyvarinen, 1999) was used to compute the ICAs. Second, the whole set of 1,391 subjects was randomly split into four folds (∼347 subjects per fold), followed by the application of the multiscale individual component clustering algorithm (MICCA) to classify ICs into *N* groups (Naveau et al., 2012). For each fold, the number of groups was automatically estimated by the algorithm (Naveau et al., 2012). The procedure was iterated seven times, leading to 28 repetitions (4-folds x 7-iterations) of the MICCA clustering. Note that for the replication sample, as the sample was not large enough to be split, we repeated single-subject ICAs 20 times for each subject, leading to 20 repetitions. Third, the *Icasso* algorithm (Himberg et al., 2004) was used to select groups of reproducible ICs (i.e., groups with ICs that were present in at least 50% of the 28 repetitions), which we identified as the “group-level components” (Salman et al., 2019). Fourth, for each group-level component, a voxel-wise *t*-score map of individual ICs was computed and thresholded using a mixture model (*P>*0.95, (Beckmann & Smith, 2004)). This process identified 56 and 58 group-level ICs for the main and BTNRH cohorts, respectively.

Then, in the main cohort, two experts (GED and MJ) reviewed all the group-level ICs and discarded ICs if their spatial map: (1) mainly covered non gray matter (i.e., CSF and WM), (2) included regions with strong signal attenuation due to susceptibility artifacts (e.g., lower frontal and lower temporal regions), (3) showed poor reproducibility (<50% of replications), and (4) were duplicates (>50% of spatial overlap across ICs (based on dice coefficients), within the main cohort). Among two duplicates, we kept the IC that was the most reproducible across the repetitions. From that selection, 30 group-level ICs were kept for further consideration.

Given that single-subject ICA in combination to MICCA leads to the presence of spatial overlap between components (Naveau et al., 2012), and because we aimed to create a brain atlas of networks without spatial overlap for easier interpretation and to be more similar to the other major adult-based atlases that also do not have any spatial overlap among networks (e.g., (Glasser et al., 2016; Yeo et al., 2011), we further removed the spatial overlap between components by attributing each voxel to a single network (among the 30 networks) based on their highest z-value.

Lastly, out of the 30 non-overlapping networks, we further excluded 6 networks that could not be identified in the BTNRH sample (i.e., <40% of overlap with any BTNRH non-overlapping components; **Supplementary Figure S3**), which led to a final number of 24 reliable and non-overlapping networks included in Dev-Atlas.

### Resting-state Network Classification and Naming

Because there is currently no consensus on how to classify and/or name a resting-state network (RSN), we chose to spatially compare each of the 24 RSNs to each of the seven networks from the Yeo Atlas (Yeo et al., 2011), computing a dice coefficient and name them by providing both anatomical and functional names (Uddin et al., 2023; Uddin et al., 2019). Each network was then classified as a subnetwork within one of the six ubiquitous large-scale functional systems (all but the limbic system as none of the networks fell into this system) based on its maximum spatial overlap.

To ensure the consistency of network classification, we also used the Network Correspondence Toolbox (NCT) (Kong et al., 2024) which provides a network classification for each network based on other major reference brain atlases (Glasser et al., 2016; Gordon et al., 2017; Ji et al., 2019; Laird et al., 2011; Power et al., 2011; Schaefer et al., 2018; Shen et al., 2013; Shirer et al., 2012) and demonstrates the level of consistency of naming and classification across all of them (**Supplementary Table S1, Supplementary Figure S5**).

### Effects of Age and Sex

Lastly, we investigated the effect of age and sex on each network’s spatial map and FNC, within the largest two datasets (PNC and HCP-D, separately). For this, we spatially reconstructed each network (using the z-map of each network as the template) within each individual, using the GIFT toolbox (through the spatial reference-based ICA approach) (https://github.com/trendscenter/gift). In GIFT, we then conducted linear model analyses in the Mancovan v1.0 toolbox to test the effect of age and sex on the spatial maps of the networks and on the FNC. Mean FD was added as a covariate of no interest. Multivariate analyses were first conducted and if significant, univariate analyses were further done. Significant effects are reported at a pFDR<0.01.

## Results

Our final main sample included 1,391 individuals (47% males, mean age=13.56 (2.7) years; PNC: n=848, PING: n=78, HCP-D: n=465; **Table 1, Supplementary Figures S1 & S2**). A total of 24 non-overlapping functional brain networks were identified as part of Dev-Atlas (**Figures 1 & 2, Supplementary Figure S4, Supplementary Table S2**). Below, we briefly describe each network within its own functional system, based on the taxonomy created by Uddin et al. (2019). Anatomical details of each network are provided in **Supplementary Table S2**.

**Figure 1:**
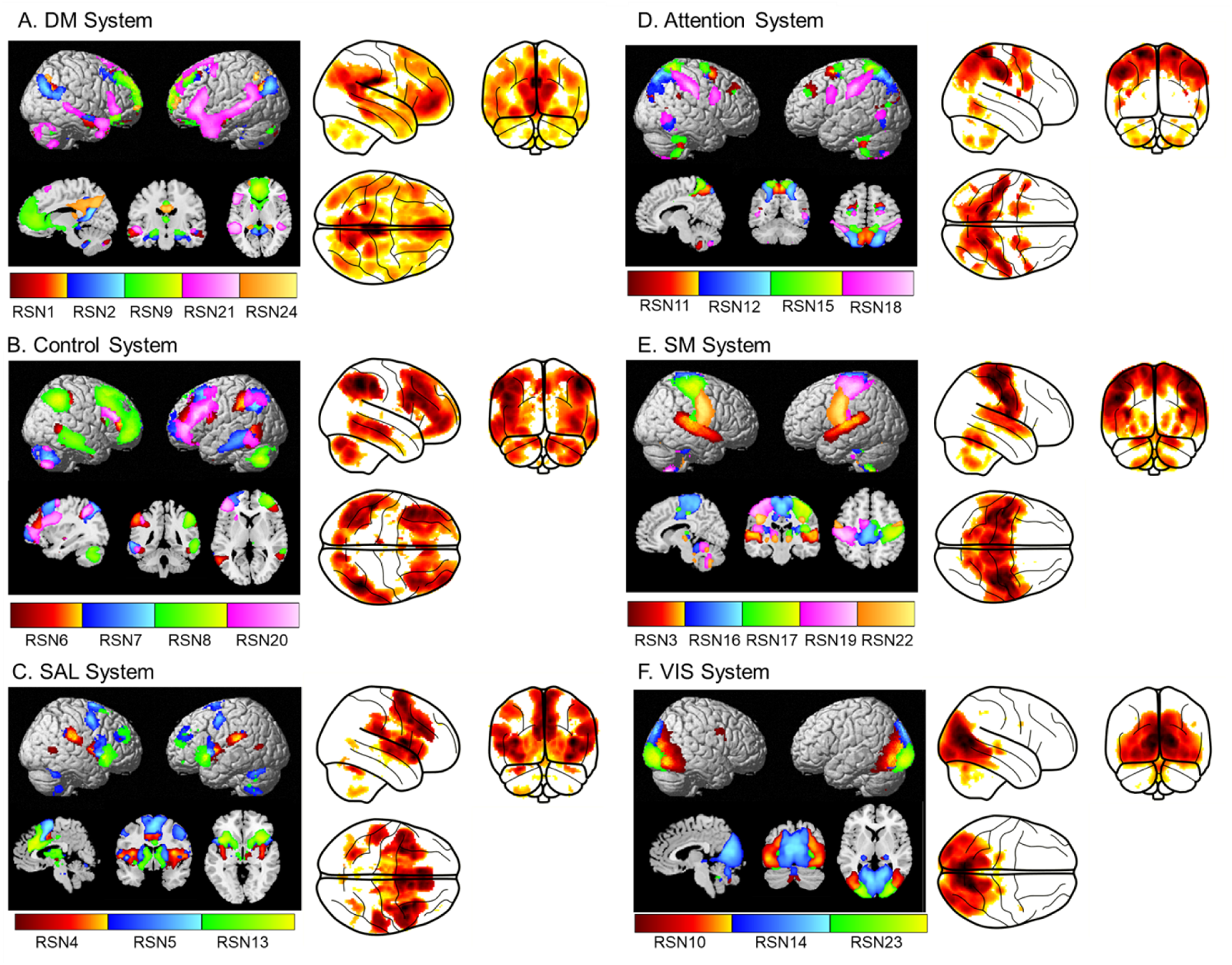
Visualization of the spatial maps of the 24 networks in Dev-Atlas, grouped by system. The right panel shows the glass brain visualization of all networks, within each system. The glass brain visualization of each network is provided in Supplementary Figure S3 and their anatomical description in Supplementary Table S2. DM= Default Mode, RSN: Resting-state network, SAL= Salience, SM=Somatomotor, VIS=Visual.

### Default mode (DM) System. Anatomical name: Medial Frontoparietal System (M-FPS). (Figure 1A, Supplementary Figure S4A)

In this system, we identified five networks. RSN01 shows the core DM network (Andrews-Hanna et al., 2010), including precuneus and medial orbitofrontal cortex, bilateral middle temporal. RSN02 covers the medial parietal and temporal cortices, such as bilateral parahippocampal gyri, which has been referred to as the medial temporal lobe (MTL) DM sub-network. RSN09 represents an anterior section of the DM network, largely encompassing anterior cingulate cortex (ACC) and medial prefrontal cortex (MPFC). RSN21 is a left-lateralized network, including left inferior frontal and middle temporal cortices, which has been described in the 17-network parcellation from Yeo et al. (2011). Lastly, RSN24 largely covers the posterior section of the DM network, with the precuneus and PCC (Doucet et al., 2011).

### Control System. Anatomical name: Lateral Frontoparietal System (L-FPS). (Figure 1B, Supplementary Figure S4B)

In this system, we identified four strongly lateralized networks, largely covering lateral parietal, frontal and temporal cortices. In detail, the RSN06 covers left-lateralized inferior frontal and parietal cortices and the posterior part of the middle temporal gyrus. The RSN07 includes left-lateralized middle frontal and temporal cortices as well as the left angular gyrus. RSN08 is right-lateralized, including middle frontal and temporal cortices, as well as the right inferior parietal cortex. Lastly, RSN20 is also a left-lateralized network, largely encompassing part of the left inferior, middle and superior frontal cortices, and to a lower level, part of the inferior parietal and temporal cortices.

### Salience (SAL) System. Anatomical name: Midcingulo-Insular System (M-CIN). (Figure 1C, Supplementary Figure S4C)

This system included three networks. RSN04 includes bilateral clusters in the insula, middle part of the superior temporal gyi and middle cingulate cortex (MCC). RSN05 covers dorsal ACC, anterior insula as well as bilateral basal ganglia (pallidum, putamen, thalamus, caudate), and bilateral cluster in the middle frontal cortex. RSN13 includes bilateral anterior insula, ACC and basal ganglia (putamen, caudate).

### Attention System. Anatomical name: Dorsal Frontoparietal System (D-FPS). (Figure 1D, Supplementary Figure S4D)

In this system, we identified four bilateral networks. RSN11 covers a large portion of the precuneus as well as bilateral clusters in the superior frontal cortex. RSN12 includes the lateral part of the precuneus, extending to the superior parietal cortex, bilaterally, as well as small posterior clusters in the inferior temporal and superior frontal cortices. RSN15 covers bilateral regions of the superior parietal cortex, and bilateral posterior regions of the superior frontal cortex. Lastly, RSN18 covers the bilateral intraparietal gyri with additional posterior clusters in the middle temporal gyri and bilateral precentral gyri.

### Somatomotor (SM) System. Anatomical name: Pericentral System (PS). (Figure 1E, Supplementary Figure S4E)

This system included five networks. RSN03 includes bilateral superior temporal gyri, typically associated with the auditory cortex. RSN16 largely covers the dorsal part of the SM network, including paracentral lobule, supplementary motor area (SMA), and pre-and postcentral gyri associated with the foot area in the homunculus. RSN17 is largely right-hemisphere lateralized, covering the right pre- and postcentral gyri associated with the left hand area in the homunculus. RSN19 is RSN17’s symmetric network, being left lateralized and covering the left pre- and postcentral gyri associated with the right hand area in the homunculus. Lastly, RSN22 largely covers the ventral part of the SM network, including bilateral pre- and postcentral gyri associated with the face area in the homunculus.

### Visual (VIS) System. Anatomical name: Occipital System (OS). (Figure 1F, Supplementary Figure S4F)

This system included three networks. RSN10 includes the lateral part of the visual network, largely covering the inferior and middle occipital cortices. RSN14 includes the medial part of the visual network, encompassing the calcarine sulcus. Lastly, RSN23 includes the most posterior part of the visual network, through the inferior occipital cortex.

**Figure 2:**
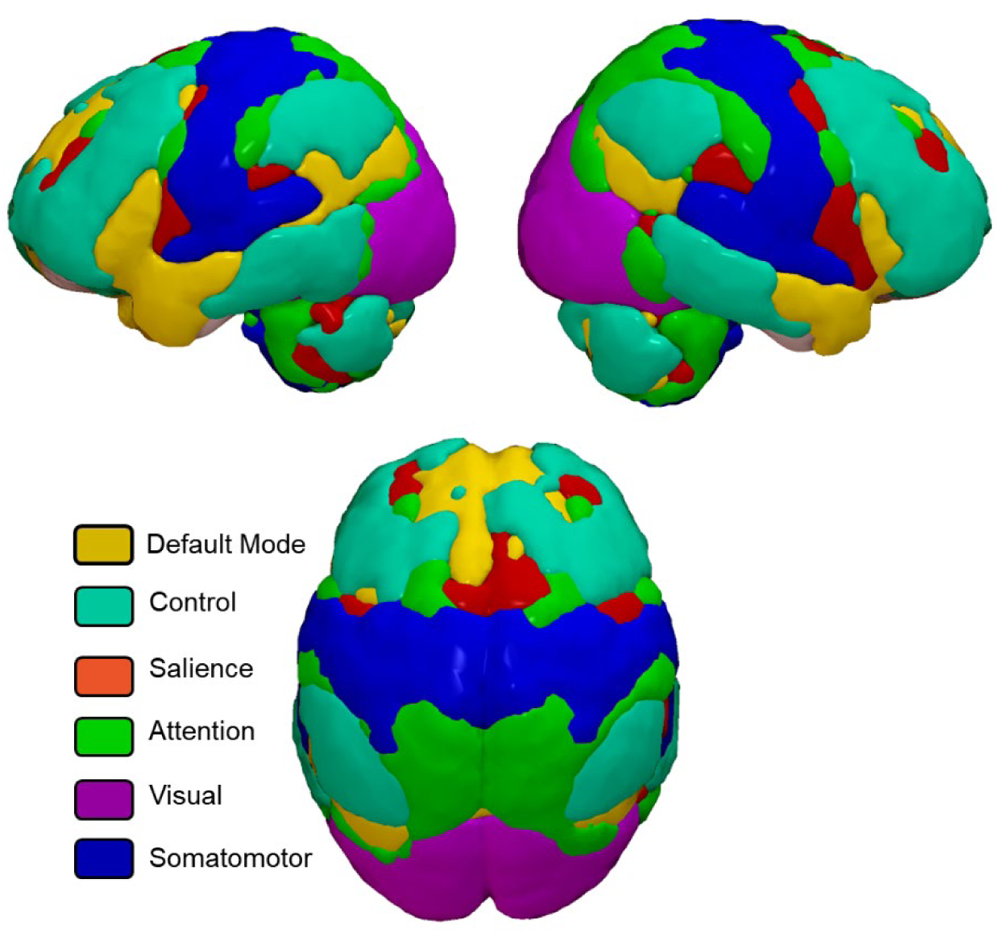
Surface-based display of the six systems from Dev-Atlas.

### Effects of Age and Sex

We further investigated the effects of age and sex on functional brain networks’ spatial maps and FNC, within the PNC and HCP-D samples separately. All results were largely similar across both samples, therefore we only report the results from the largest sample (PNC) below, while the results related to the HCP-D are provided in Supplementary Material (**Supplementary Figures S6 & S7**). There was no significant sex-by-age interaction for any of the models tested, so we only focus on the main effects.

### Spatial maps

*Effects of age*: Multivariate analyses revealed a main effect of age, both positive and negative, on most of the networks in the atlas, except the networks from the VIS system (16 / 24 networks). The major effects were detected in the DM, Control and SAL systems (**Figure 3, Supplementary Table S3**). For simplicity, we discuss the effects of age at the system level, although each network’s detail is provided in **Supplementary Table S3**. In the DM system, older age was associated with higher spatial integrity of regions largely located in the medial and lateral prefrontal cortex, and with lower integrity in regions mostly located in the posterior part of the network, namely medial and lateral parietal lobe, medial temporal lobe, and lateral temporal cortex. In the Control system, older age was associated with higher spatial integrity of the lateral frontal regions and with lower integrity of the posterior regions, including posterior temporal and parietal regions. In the SAL system, older age was associated with higher spatial integrity of the ACC and subcortical regions, and with lower integrity with insula and superior temporal cortices. In the other systems, the effects were more limited and widespread across the parietal and temporal lobes.

**Figure 3:**
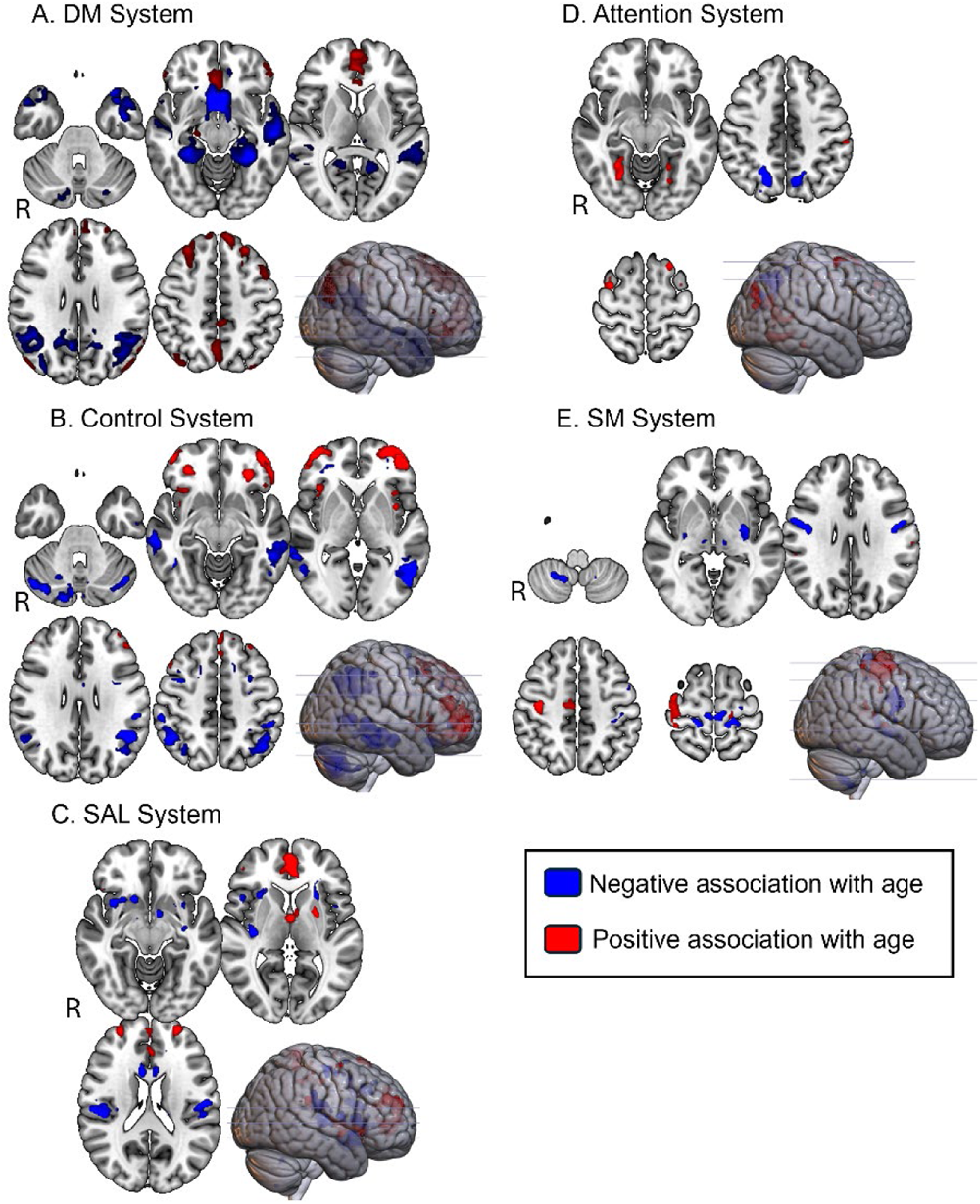
Effect of age on the spatial maps of each Dev-Atlas network, organized by system. Blue and red regions display negative and positive associations with age, respectively. Views are in radiological orientation (Right side shows the left hemisphere).

*Effects of sex*: We found relatively small sex differences in 13 networks of the atlas, with the networks from the Control and DM systems the most impacted (**Figure 4, Supplementary Table S4**). In the DM system, males showed temporal and left frontal regions more strongly integrated than females. In contrast, the core medial and ventral regions of the DM network (precuneus, orbitofrontal cortex, medial temporal lobe) were more strongly integrated in females than males. In the Control system, the parietal regions were stronger in the males than the females, while clusters in the lateral prefrontal cortex were more strongly integrated in the female than in the male participants. In the SAL system, males showed greater integrity in the anterior insula, while for females it was in the subcortical regions. The other sex effects related to the other systems were much smaller in size and intensity and are detailed in **Supplementary Table S4**.

**Figure 4:**
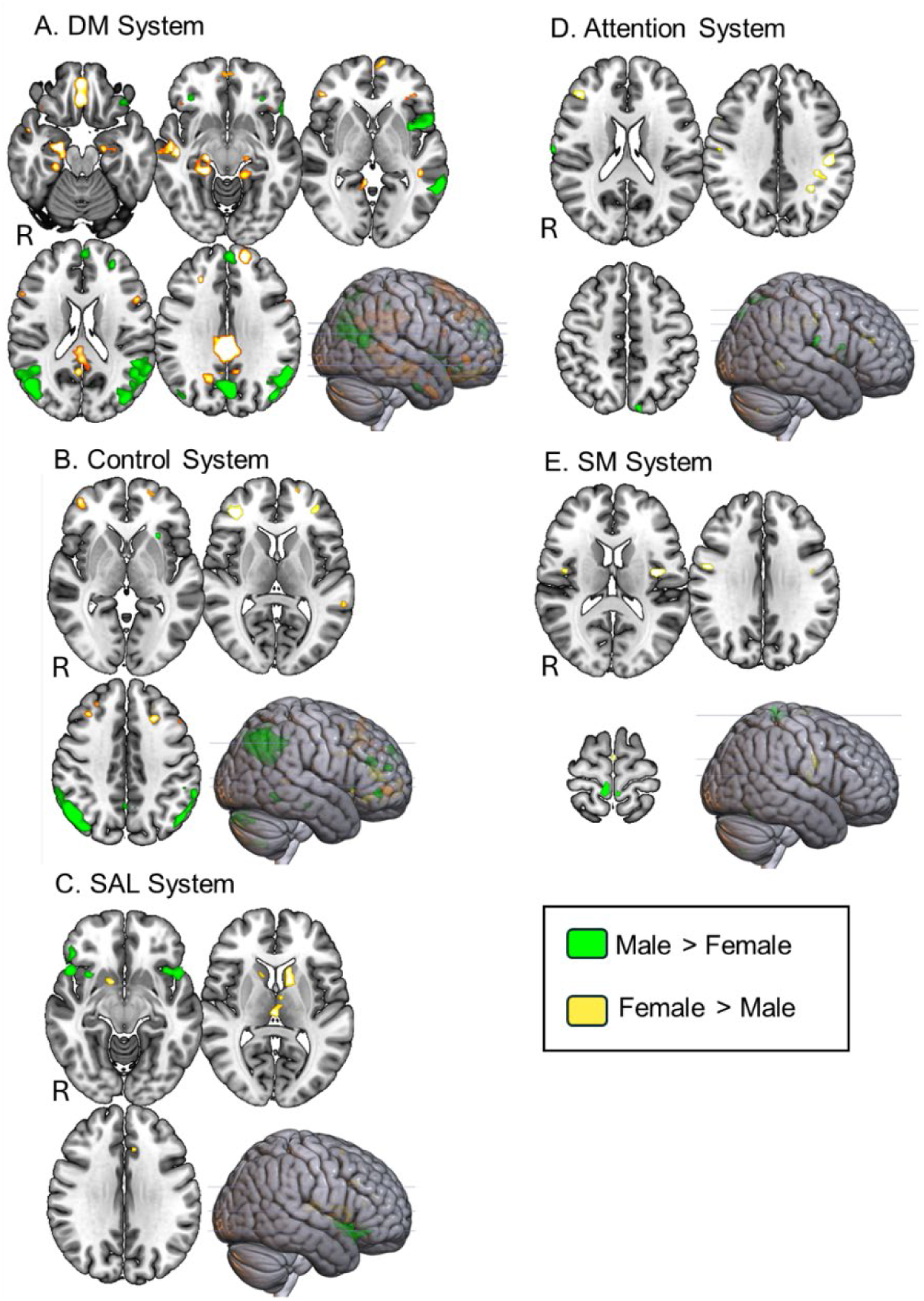
Effect of sex on the spatial maps of each Dev-Atlas network, organized by system. Green and yellow regions display male > female and female > male, respectively. Views are in radiological orientation (Right side shows the left hemisphere).

### FNC

*Effects of age*: Multivariate analyses revealed a main effect of age on FNC across all networks. We therefore conducted further univariate analyses. Across networks, there was a clear pattern in which older age was associated with reduced FNC between the networks from the DM or the Control System and the networks from the other systems (**Figure 5A**). Additionally, older age was associated with increased FNC within most of each system. In particular, greater FNC was revealed among all networks within the SM, as well as between the Attention and SAL systems, with increasing age.

*Effects of sex*: To a lesser degree, we revealed an effect of sex on FNC across some networks (**Figure 5B**). The main differences appeared within the DM, Attention and SAL systems. Males showed weaker FNC within the DM system and between the Attention and SAL system, while greater FNC within the Attention system, compared with females.

**Figure 5:**
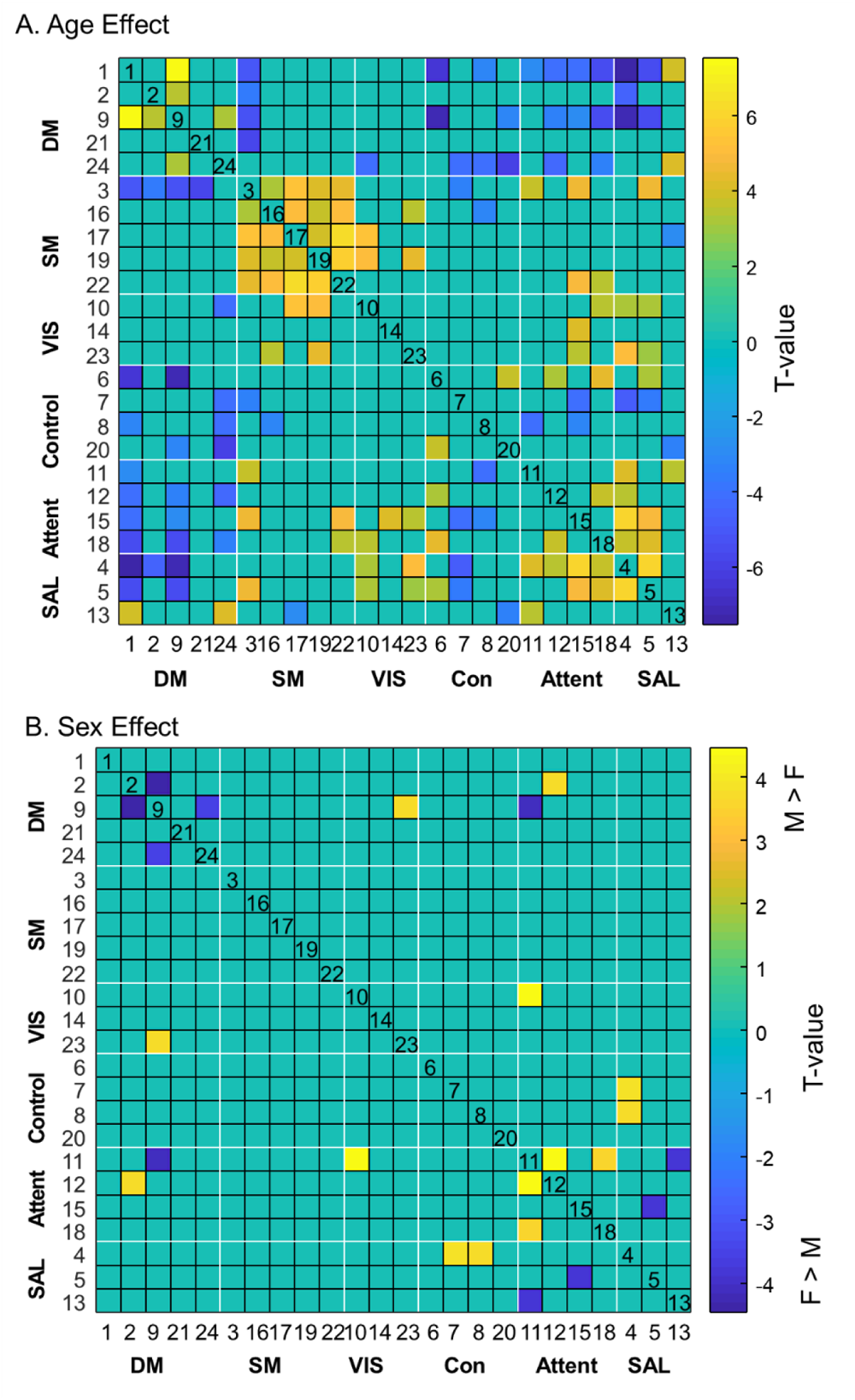
Significant effect of age and sex on the FNCs. Results of the univariate analyses are displayed. The significant threshold was set up at pFDR<0.01.

### Data Sharing

The thresholded binary and non-binary spatial maps of each of the 24 networks included in Dev-Atlas are freely available at https://github.com/BRAICLab/DevAtlas. Given the impact of age, we also computed two additional sub-atlases after splitting the whole sample based on the median age (13.83 years old) into a younger (8.0-13.82) and older (13.83-17.9) groups. Following the same analysis pipeline, 20 and 22 reliable networks were then identified in the younger (Dev-AtlasA) and older (Dev-AtlasB) groups, respectively. These networks are also freely available for more age-specific analyses.

## Discussion

This study introduces a novel reference atlas of functional brain networks for typically developing adolescents, named Dev-Atlas. To construct this atlas, we used datasets from three large independent studies (PNC, HCP-D and PING), as well as a smaller dataset locally collected from typically developing children and adolescents aged 8 to 17 years-old. To further characterize the Atlas, we investigated the impact of age and sex on the spatial topography and functional connectivity of each identified network. Consistent with published developmental literature (Cai et al., 2018; Di Martino et al., 2014; Dosenbach et al., 2010; Ernst et al., 2015; Gu et al., 2015; Sole-Padulles et al., 2016; Uddin et al., 2010; Uddin et al., 2011), we reported both positive and negative robust effects of age on a large majority of the networks, at both the spatial topography and functional connectivity level, while the observed sex differences were more limited.

This new atlas was constructed using 1,391 rsfMRI datasets and includes 24 reliable networks, classified within six functional systems: Default-Mode, Control, Salience, Attention, Somatomotor and Visual, based on their spatial similarity with the corresponding system defined by Yeo et al. (2011). While there is no clear consensus on the spatial definition of each brain system and their functional naming, we chose to use one of the most largely used, based on Yeo et al. (2011) as it was built based on 1,000 participants and has been found as reliable (Doucet et al., 2019). However, we also did a secondary analysis using the open-access Network Correspondence Toolbox (NCT) (Kong et al., 2024), and were able to confirm the classification of each network among other brain atlases with high reliability (**Supplementary Table S1, Supplementary Figure S5**). As expected, all networks were relatively spatially similar to what has been described in adult populations (Fu et al., 2024; Yeo et al., 2011), with some key differences. In Dev-Atlas, most of the functional brain systems included a larger number of networks than typically reported in adult functional brain parcellations (e.g., (Yeo et al., 2011)) which is consistent with the fact that early life is associated with greater brain parcellation (Fu et al., 2024; Gu et al., 2015).

In comparison to adult-derived functional brain networks, the most significant differences could be noticed in the significant impacts of age at both spatial topography and functional connectivity levels. Most of age impacts in adult functional brain networks have typically been largely negative (Andrews-Hanna et al., 2007; Damoiseaux, 2017; Doucet et al., 2020); while in Dev-Atlas, a relatively equal number of positive and negative changes can be noticed. These changes are likely to reflect functional maturation of the brain networks throughout typical development. Accordingly, as age increases, networks within a specific system, especially in the SM System, were found to become more connected. In early life, the SM system has been described as a major hub, and becomes less central as children grow, and by adulthood, the SM system is typically not considered a central hub anymore (Oldham & Fornito, 2019). In Dev-Atlas, the SM system includes five networks, with each covering specific sections of the somatomotor cortex. The high positive FC with increasing age may likely lead to lower parcellation of this system in adulthood (i.e., less networks as typically reported in other adult brain parcellations). In contrast, we also revealed that increasing age was associated with negative FC between the DM networks and networks from all the other systems as well as between the Control system and networks from both the Attentional and SAL systems. These negative links may likely reflect greater functional specialization and segregation of both DM and Control systems as adolescents become adults. Indeed, the DM network is known to become one of the most central brain networks in adult populations (Buckner et al., 2008; Buckner & DiNicola, 2019; Raichle, 2015), supporting self-referential processing that are typically still developing until mid-adulthood (Andrews-Hanna et al., 2014). Our findings are also in line with seminal works that have demonstrated the increasing specialization of the DM through both increased within-network FC (Fair et al., 2008; Fan et al., 2021) and decreased FC with other networks throughout development (Dosenbach et al., 2010). Similarly, the Control, Attentional and SAL systems are known to work together to support higher-order cognitive functions and externally-oriented attention (Buckner et al., 2013; Doucet et al., 2011; Smith et al., 2009; Uddin et al., 2019). Throughout development, our findings point to a refinement of the Control system, through a better delineation from the default, attention and salience systems. These finding broadly align with report increases in network segregation across the lifespan (Vij et al., 2018).

At the spatial level, we revealed interesting pattern of spatial changes within networks as age increases, which is consistent with the previous work by Sole-Padulles et al. (2016) that was conducted on 113 typically-developing youth aged 7-18 years. In particular, we revealed an overall positive association with anterior DM and a negative association with the posterior DM networks (Sole-Padulles et al., 2016). This anterior/ posterior segregation was also revealed to some level for the Control and SAL networks and may reflect the late maturation of the frontal lobe (Walhovd et al., 2017). Within the SAL system, we also identified a large impact of age, including the right but not so much the left insula (RSN04 vs RSN13, **Supplementary Table S3**), which is consistent with previous findings suggesting a strong developmental impact on the laterality of the fronto-insular cortex (Uddin et al., 2011). In fact, it is likely that the right front-insular cortex undergoes critical maturation with age, providing access to attentional and working memory resources (Uddin et al., 2011).

In contrast to the large reproducible effect of age across the brain networks, we failed to reveal any statistically significant age-by-sex interaction and only reported minimal sex effects, which is largely consistent with previous rsfMRI studies reporting none or small sex effects in youth (Satterthwaite et al., 2015; Sole-Padulles et al., 2016). Although sex differences are evident throughout development at the pubertal (Herting & Sowell, 2017) and structural brain (Gur & Gur, 2016; Modabbernia et al., 2021; Wierenga et al., 2020) levels; sex differences in fMRI have not been as consistently reported or are typically relatively weak in children and adolescent samples (Rubia, 2013; Satterthwaite et al., 2015; Sole-Padulles et al., 2016). In our data, the largest sex-related spatial differences were revealed within the spatial topography of the DM system, and particularly within the RSN 21, which includes left-lateralized frontal-parietal regions and has been commonly linked to language processing (Doucet et al., 2011), with males showing stronger spatial integrity than females. This finding is in line with the study by Xu et al. (2020) that described sex differences on brain effective connectivity in the language network. Combined with our finding, such results provide further implication on the origins of the sex differences revealed in children diagnosed with language disorders, such as dyslexia (Rutter et al., 2004), stuttering (Drayna et al., 1999) or developmental language disorder (Chilosi et al., 2023), which seem to affect males more often than females.

Other sex differences were revealed in term of between-network FC with males showing greater between system FC than females, while females seem to show higher within-network FC within the DM system than males. Consistently, Sattertwaite et al. (2015) reported that males have a tendency to have greater between-network connectivity than females, although this effect was small (d=0.33) but stable across the adolescent period, as they also did not detect an age-by-sex interaction. However, literature remains unclear on that aspect as some studies have reported strong puberty-by-sex interaction in the DM network functional connectivity in adolescents (Ernst et al., 2019). However, their sample focused on 13-15 year olds and it is therefore possible that such interaction may only be identified in very limited age groups. It will be important to further investigate sex effects in groups of adolescents with more limited age ranges.

Although Dev-Atlas offers a realistic option for standardizing the definition of functional brain networks in youth, we acknowledge its limitations. First, the age range from 8-17 years can be viewed as arbitrary and too extensive given the large structural and functional brain changes occurring during this age period (Uddin et al., 2011); however, there is currently no exact age that starts or ends puberty on each child, especially as the pubertal onset differs between girls and boys (Hoyt et al., 2020). We therefore followed the most common age criterion across developmental literature (Somerville et al., 2018). In order to mitigate effects of the broad age range, we also provide the spatial map of the networks identified in two subgroups of participants, equally split based on the median age to provide more specific age-specific networks. Second, Dev-Atlas was created using a combination of single-subject ICA and spatial clustering following the validated approach by Naveau et al. (2012); which was further used to create Atlas55+, a reference brain atlas for late adulthood (Doucet et al., 2020). We chose this volume-based approach, in comparison to a surface-based approach as applied in the brain atlas by Yeo et al. (2011). The major reason for this was to include cortical, subcortical and cerebellar regions in this atlas. Indeed, surface-based atlases include cortical regions only. However, during adolescence, both subcortical and cerebellar regions go through significant changes, in term of both structure (Goddings et al., 2014; Tiemeier et al., 2010) and FC with cortical networks (Chung et al., 2020; Gaiser et al., 2024; Kundu et al., 2018; Sathyanesan et al., 2019). As reported in Supplementary Table S2, the large majority of the networks we identified include some cerebellar and/or subcortical regions, which reinforces our choice of a volume-based approach. While ICA is a well-known and well-validated data-driven approach wildly used in neuroscience to extract meaningful imaging features (Calhoun et al., 2009; Fu et al., 2024; Liu et al., 2009), we cannot exclude the possibility that the clustering approach did not influence the current findings (Abou Elseoud et al., 2011). Third, age was only investigated using a linear approach in these cross-sectional datasets. It is therefore possible that we have only captured a portion of developmental changes in the functional brain networks, but this aspect was not a direct scope of the study. It will be important to further investigate the brain networks and the impact of age throughout adolescence, in a longitudinal setting such as using the ABCD dataset when the longitudinal collection will be fully completed (Casey et al., 2018). Lastly, this study specifically focused on the spatial characteristics of typically developing functional brain networks; other characteristics such as their dynamics or the association with mental health in adolescents were not. In particular, it will be important to investigate the spatial definition of the Dev-Atlas networks as a function of increasing risk for psychopathology.

## Conclusion

We built a new reference brain atlas composed of 24 reliable networks for typical adolescence. We hope that Dev-Atlas will be able to be used to capture adolescent-specific functional features which might boost accuracy in mental health research and advance our understanding of this age group and its increased risk to psychopathology. We further provide information on both age and sex impacts on the spatial topography and functional connectivity of each identified network for a better network characterization. This atlas is freely available and is also available as part of the GIFT toolbox (http://trendscenter.org/software/gift).

