## Supplementary Material for "Dev-Atlas: A reference atlas of functional brain networks for typically developing adolescents"

### Rs-fMRI Preprocessing

For each individual's rsfMRI run, the following preprocessing was performed. First, a reference volume and its skull-stripped version were generated using a custom methodology of *fMRIPrep*. Head-motion parameters with respect to the BOLD reference (transformation matrices, and six corresponding rotation and translation parameters) are estimated before any spatiotemporal filtering using MCFLIRT (FSL 6.0.5.1:57b01774, Jenkinson et al. 2002). BOLD runs were slice-time corrected to 1.47s (0.5 of slice acquisition range 0s-2.94s) using 3dTshift from AFNI (Cox and Hyde 1997, RRID:SCR_005927). The BOLD time-series (including slice-timing correction when applied) were resampled onto their original, native space by applying the transforms to correct for head motion. These resampled BOLD time-series will be referred to as *preprocessed BOLD in original space*, or *preprocessed BOLD*. The BOLD reference was then co-registered to the T1w reference using bbregister (FreeSurfer) which implements boundary-based registration (Greve and Fischl 2009). Co-registration was configured with six degrees of freedom. Several confounding time-series were calculated based on the *preprocessed BOLD*: framewise displacement (FD), DVARS, and three region-wise signals. FD was computed using two formulations following Power (absolute sum of relative motions, Power et al. (2014)) and Jenkinson (relative root mean square displacement between affines, Jenkinson et al. (2002)). FD and DVARS are calculated for each functional run, both using their implementations in *Nipype* (following the definitions by Power et al. 2014). The three signals are extracted within the CSF, the WM, and the whole-brain masks. Additionally, a set of physiological regressors were extracted to allow for component-based noise correction (*CompCor*, Behzadi et al. 2007). Principal components are estimated after high-pass filtering the *preprocessed BOLD* time-series (using a discrete cosine filter with 128s cut-off) for the two *CompCor* variants: temporal (tCompCor) and anatomical (aCompCor). tCompCor components are then calculated from the top 2% variable voxels within the brain mask. For aCompCor, three probabilistic masks (CSF, WM and combined CSF+WM) are generated in anatomical space. The implementation differs from that of Behzadi et al. in that instead of eroding the masks by 2 pixels on BOLD space, a mask of pixels that likely contain a volume fraction of GM is subtracted from the aCompCor masks. This mask is obtained by dilating a GM mask extracted from the FreeSurfer's *aseg* segmentation, and it ensures components are not extracted from voxels containing a minimal fraction of GM. Finally, these masks are resampled into BOLD space and binarized by thresholding at 0.99 (as in the original implementation). Components are also calculated separately within the WM and CSF masks. For each CompCor decomposition, the *k* components with the largest singular values are retained, such that the retained components' time series are sufficient to explain 50 percent of variance across the nuisance mask (CSF, WM, combined, or temporal). The remaining components are not considered further. The head motion estimates calculated in the correction step were also placed within the corresponding confounds file. The confound time series derived from head motion estimates and CSF, WM, and whole-brain signals were expanded with the inclusion of temporal derivatives and quadratic terms for each (Satterthwaite et al. 2013). Frames that exceeded a threshold of 0.5 mm FD or 1.5 standardized DVARS were annotated as motion outliers. Additional nuisance timeseries are calculated by means of principal components analysis of the signal found within a thin band (*crown*) of voxels around the edge of the brain, as proposed by (Patriat, Reynolds, and Birn 2017). The BOLD time-series were resampled into standard space, generating a *preprocessed BOLD run in MNI152NLin2009cAsym space*. First, a reference volume and its skull-stripped version were generated using a custom methodology of *fMRIPrep*. Automatic removal of motion artifacts using independent component analysis (ICA-AROMA, Pruim et al. 2015) was performed on the *preprocessed BOLD on MNI space* time-series after removal of non-steady state volumes and spatial smoothing with an isotropic, Gaussian kernel of 6mm FWHM (full-width half-maximum). Corresponding "non-aggressively" denoised runs were produced after such smoothing. Additionally, the "aggressive" noise-regressors were collected and placed in the corresponding confounds file. All resamplings can be performed with *a single interpolation step* by composing all the pertinent transformations (i.e. head motion transform matrices, susceptibility distortion correction when available, and co-registrations to anatomical and output spaces). Gridded (volumetric) resamplings were performed using antsApplyTransforms (ANTs), configured with Lanczos interpolation to minimize the smoothing effects of other kernels (Lanczos 1964). Non-gridded (surface) resamplings were performed using mri_vol2surf (FreeSurfer).

Following this, we further applied denoising strategies using the nilearn package, using the ica_aroma strategy (https://nilearn.github.io/stable/modules/generated/nilearn.interfaces.fmriprep.load_confounds_strategy.html#nilearn.interfaces.fmriprep.load_confounds_strategy), which included high pass filtering, basic white matter and CSF correction, global signal regression, and regression of the confounds derived from ICA-AROMA. This pipeline was chosen based on the recommendations by Ciric et al. (2017) showing that this approach is among the best to prevent the impact of head motion on functional connectivity.

**<u>Description of the sample</u>**

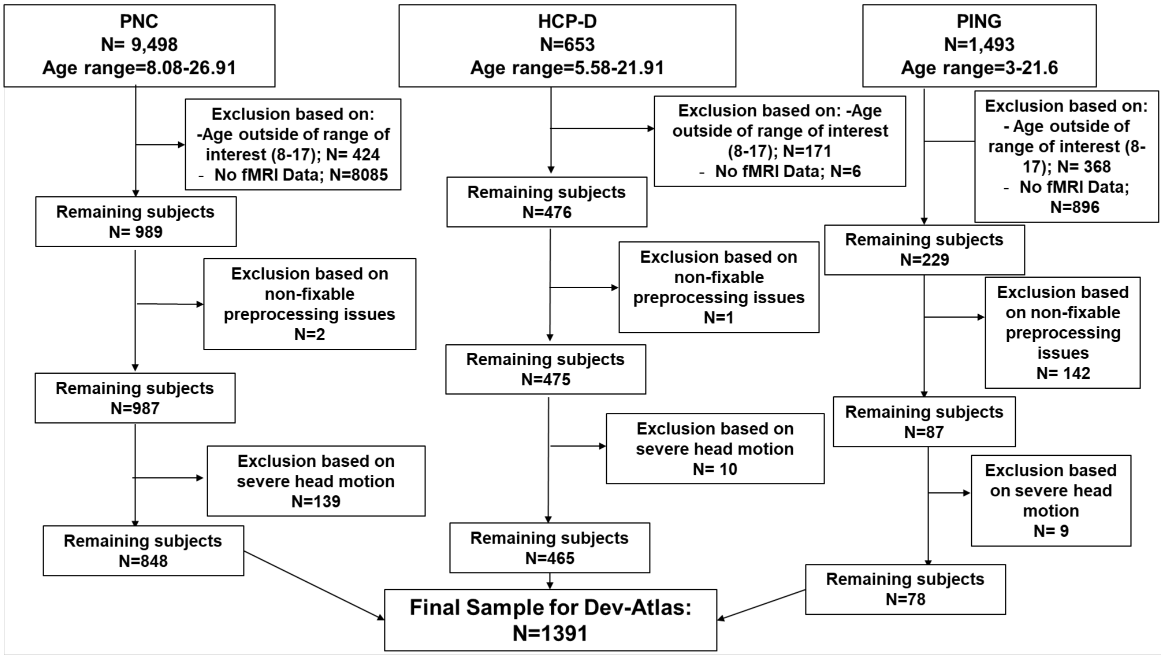

**Supplementary Figure S1: Description of the pipeline leading to the final sample.**

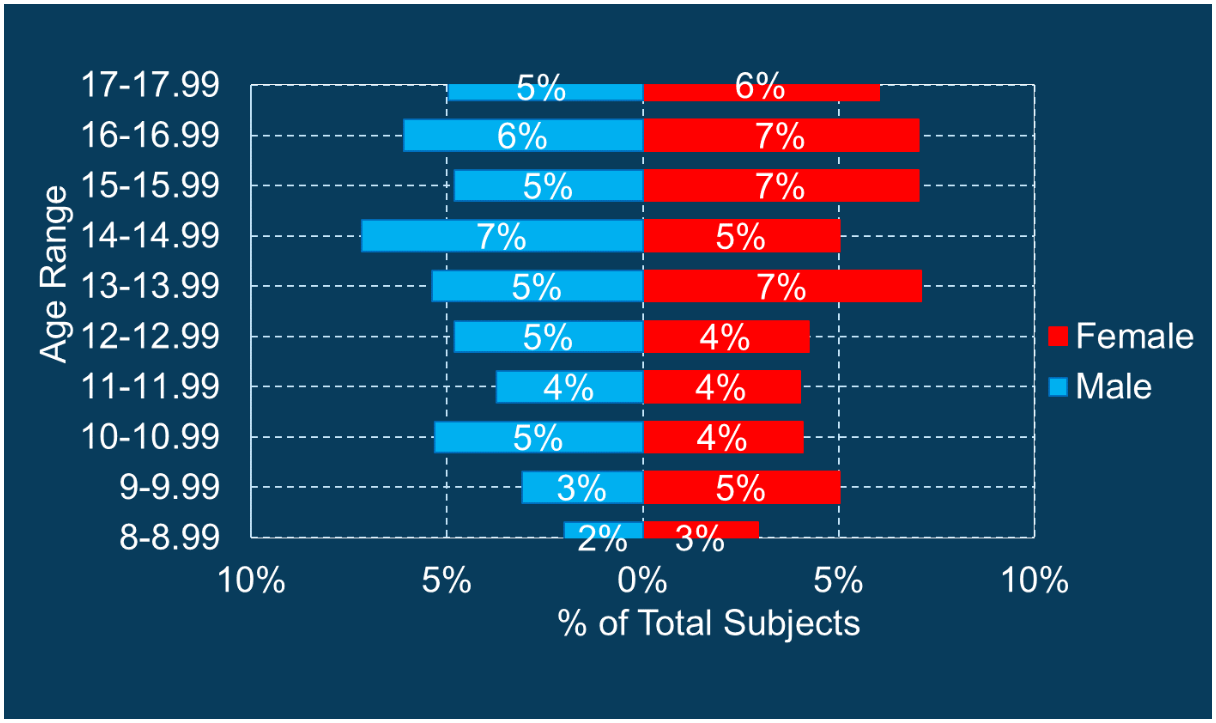

**Supplementary Figure S2: Sex distribution across the 1391 participants included.**

**Outputs of MICCA**

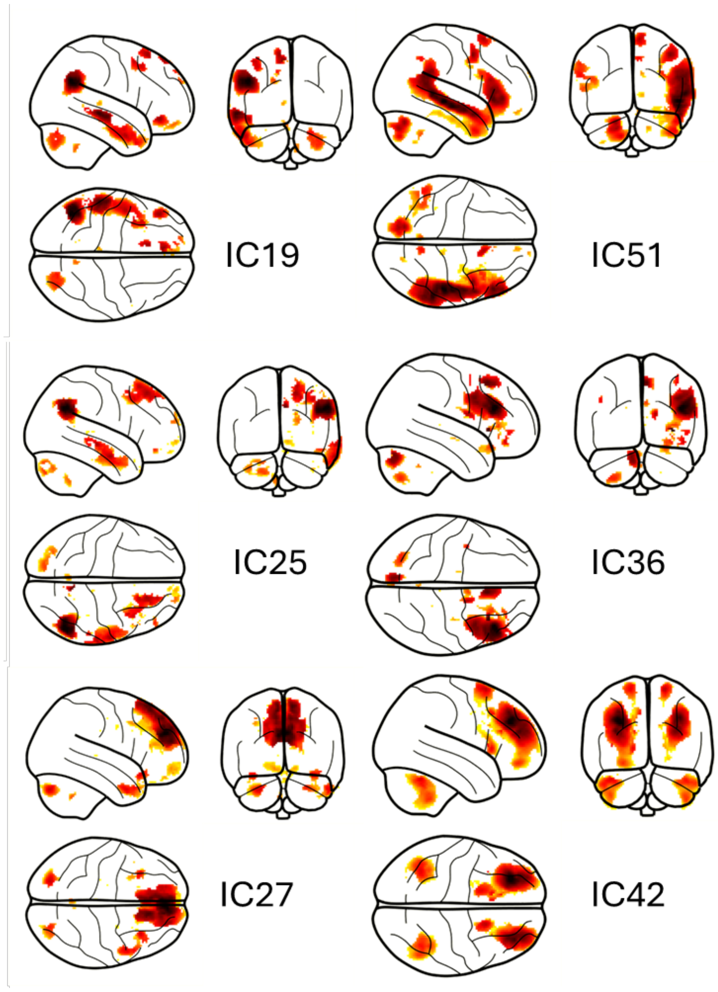

**Supplementary Figure S3: Glass-brain visualization of the independent components (ICs) not found in the BTNRH sample and therefore excluded from Dev-Atlas.**

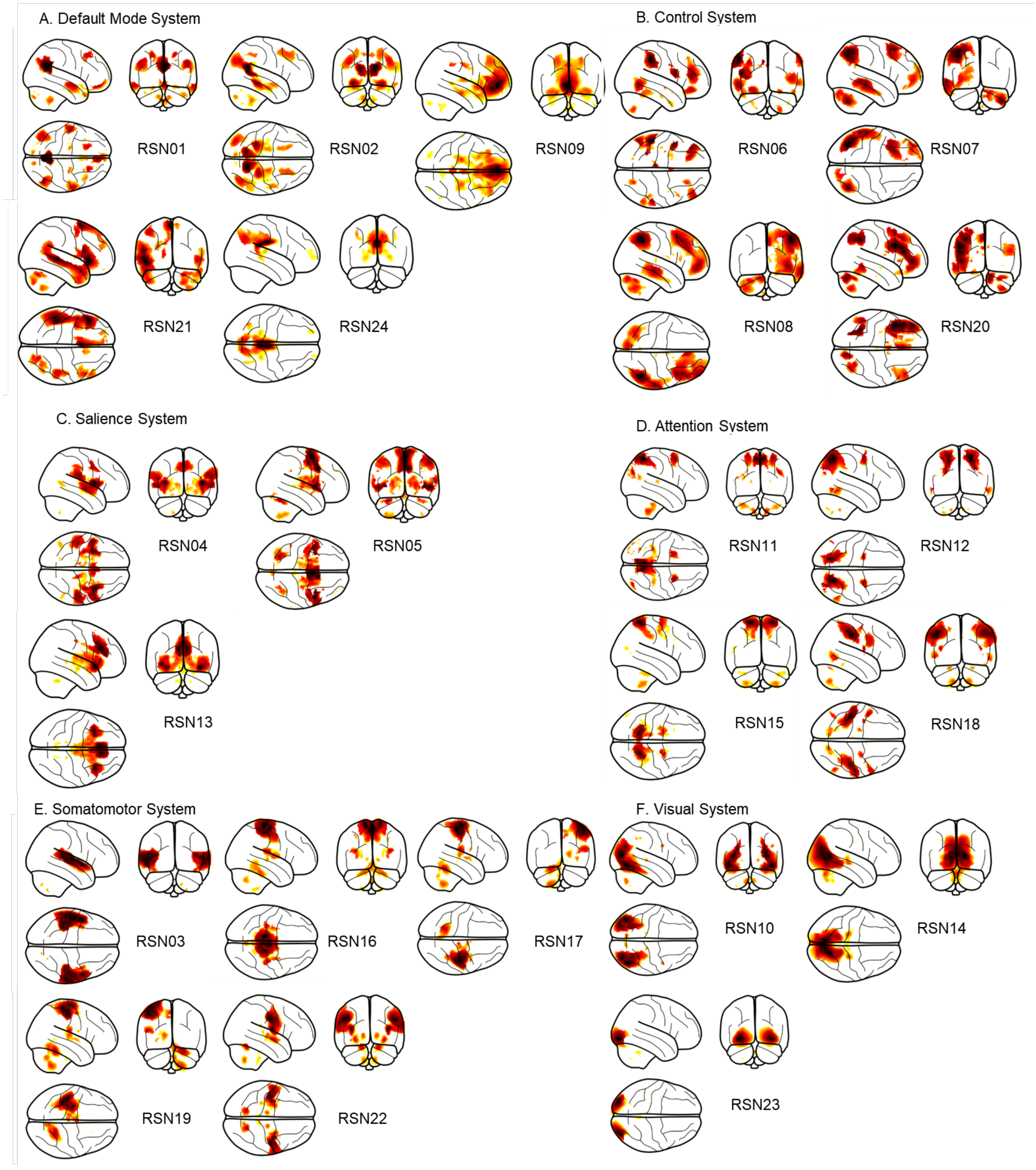

**Supplementary Figure S4: Glass brain visualization of each of the 24 networks included in Dev-Atlas**

**Supplementary Table S1: Network attribution based on the largest spatial overlap between each network from Dev-Atlas and 7 other brain atlases, based on the Network Correspondence Toolbox.**

| RSN ID | Dev-Atlas | Yeo2011 (7) | Yeo2011 (17) | Laird2011 | Power2011; Gordon2017 | Shirer2012 | Shen2013 | Glasser2016; Ji2019 | Schaefer2018; Kong2021 |
|---|---|---|---|---|---|---|---|---|---|
| 1 | **DMN** | Default | DefaultA | DivergentCog1 | Default | DorsalDMN | Default | Default | DefaultA |
| 2 | **DMN** | Default | DefaultC | DivergentCog1 | Context | VentralDMN | Default | Default | DefaultC |
| 9 | **DMN** | Default | DefaultA | Emo/Interoception2 | Default | DorsalDMN | Default | Default | DefaultA |
| 21 | **DMN** | Default | DefaultB | Emo/Interoception1 | Language | Language | MedFront | Language | Language |
| 24 | **DMN** | Control | ControlC | DivergentCog1 | ParMemory | Precuneus | Default | FrontPar | ControlC |
| 6 | **Control** | Control | ControlA | DivergentCog3 | FrontPar | PostSal | FrontPar | FrontPar | Sal/VenAttnB |
| 7 | **Control** | Control | ControlB | DivergentCog3 | Default | RECN | FrontPar | FrontPar | ControlB |
| 8 | **Control** | Control | ControlB | DivergentCog6 | FrontPar | LECN | FrontPar | FrontPar | ControlB |
| 20 | **Control** | Control | ControlA | DivergentCog3 | FrontPar | RECN | FrontPar | FrontPar | ControlA |
| 11 | **Attention** | DorsAttn | ControlC | Mot/Visspatial 2 | DorsAttn | VentralDMN | SalSubcor | PostMulti | DefaultC |
| 12 | **Attention** | DorsAttn | DorsAttnA | Mot/Visspatial 2 | DorsAttn | Visuospatial | VisAssoc | DorsAttn | DorsAttnA |
| 15 | **Attention** | DorsAttn | DorsAttnB | Mot/Visspatial 4 | Premotor | VentralDMN | VisAssoc | Somatomotor | DorsAttnB |
| 18 | **Attention** | DorsAttn | DorsAttnB | Mot/Visspatial 3 | Premotor | Visuospatial | Motor | DorsAttn | DorsAttnB |
| 4 | **SAL** | Sal/VenAttn | Sal/VenAttnA | DivergentCog5 | CingOperc | PostSal | Motor | CingOperc | Sal/VenAttnA |

| | | | | | | | | | |
|---|---|---|---|---|---|---|---|---|---|
| 5 | **SAL** | Sal/VenAttn | Sal/VenAttnA | Mot/Visspatial 1 | CingOperc | AntSal | SalSubcor | CingOperc | Sal/VenAttnA |
| 13 | **SAL** | Sal/VenAttn | Sal/VenAttnB | Emo/Interoception4 | Salience | AntSal | SalSubcor | CingOperc | ControlC |
| 3 | **SMN** | Somatomotor | SomatomotorB | DivergentCog4 | Auditory | Auditory | Motor | Auditory | Auditory |
| 16 | **SMN** | Somatomotor | SomatomotorA | Mot/Visspatial 4 | FootSM | Sensorimotor | Motor | Somatomotor | SomatomotorA |
| 17 | **SMN** | Somatomotor | SomatomotorA | Mot/Visspatial 3 | HandSM | Sensorimotor | Motor | Somatomotor | SomatomotorA |
| 19 | **SMN** | Somatomotor | SomatomotorA | Mot/Visspatial 3 | HandSM | Sensorimotor | Motor | Somatomotor | SomatomotorA |
| 22 | **SMN** | Somatomotor | SomatomotorB | DivergentCog5 | FaceSM | Sensorimotor | Motor | Somatomotor | SomatomotorB |
| 10 | **VIS** | Visual | VisualA | Visual1 | LatVis | HighVisual | VisAssoc | Visual2 | VisualA |
| 14 | **VIS** | Visual | VisualB | Visual3 | MedVis | PrimVisual | VisualA | Visual1 | VisualB |
| 23 | **VIS** | Visual | VisualA | Visual2 | LatVis | HighVisual | VisualB | Visual1 | VisualC |

**Supplementary Figure S5: Example of the network concordance for RSN 01 with networks from other brain atlases.** Values represent dice coefficients between RSN01 and each network of a corresponding atlas, based on the NCT.

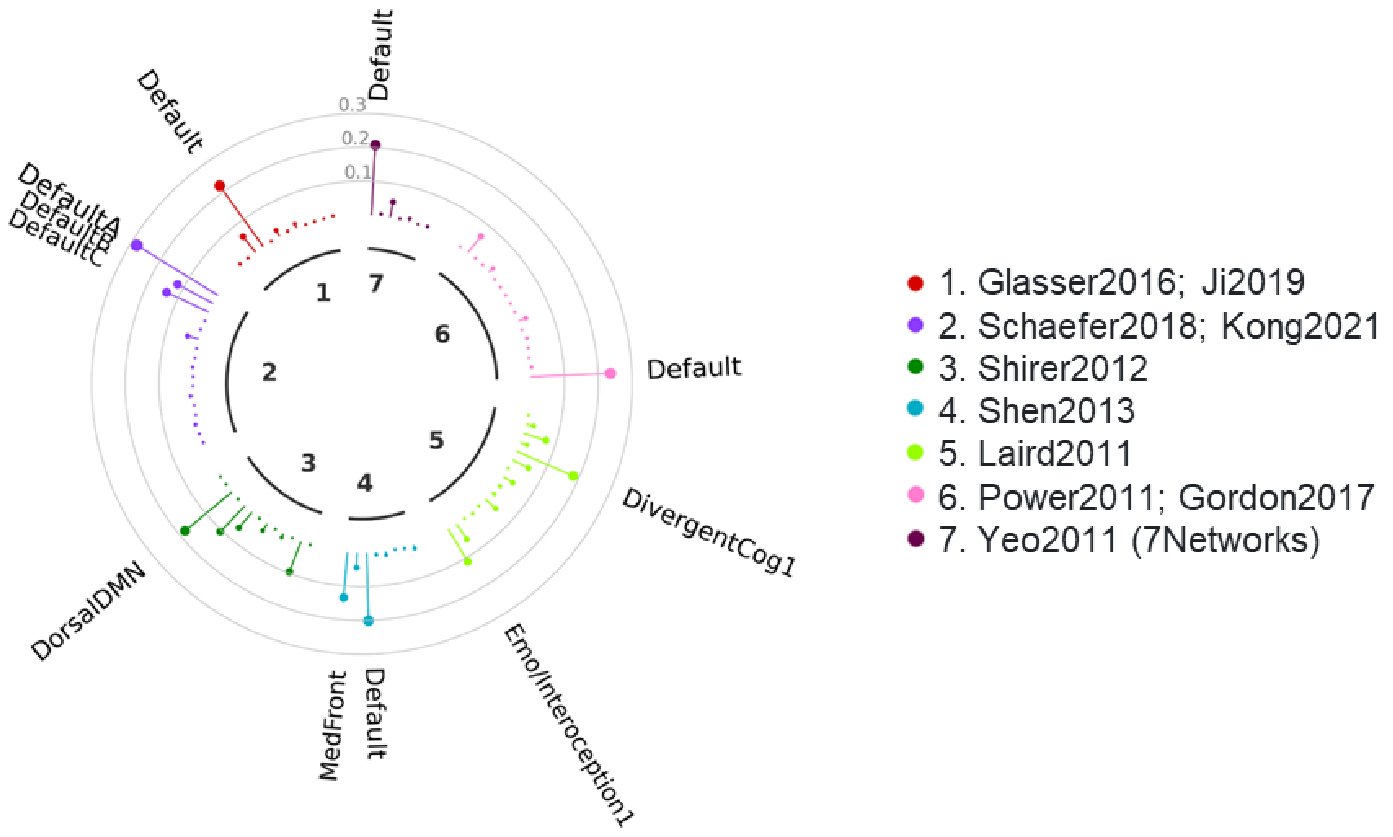

**Supplementary Table S2: Anatomical Description of the 24 Functional Brain Networks included in Dev-Atlas.**

| | Brain Region | Peak | MNI coordinate (x,y,z) | Volume (mm3) |
|---|---|---|---|---|
| **DM System** | | | | |
| RSN 01 | Precuneus_R | 12.4 | 2, -62, 26 | 8109 |
| | Angular_R | 10.3 | 48, -64, 30 | 2340 |
| | Med_Orb_Frontal_R | 10 | 5, 49, -14 | 698 |
| | Temporal_Mid_L | 9.8 | -60, -8, -14 | 1892 |
| | Temporal_Mid_R | 9.2 | 60, -4, -16 | 1500 |
| | Angular_L | 9 | -48, -66, 28 | 2983 |
| | Frontal_Mid_L | 9 | -23, 28, 48 | 814 |
| | Frontal_Mid_R | 8.5 | 24, 28, 48 | 389 |
| | Cerebellum_9_R | 6.7 | 6, -50, -46 | 46 |
| RSN 02 | Precuneus_R | 16.4 | 16, -56, 22 | 9455 |
| | Parahippocampus_L | 14.2 | -30, -38, -14 | 2667 |
| | Parahippocampus_R | 13.2 | 28, -34, -14 | 2014 |
| | Occipital_Mid_L | 11.4 | -37, -80, 30 | 2443 |
| | Occipital_Mid_R | 11.2 | 44, -72, 30 | 2178 |
| | Frontal_Sup_R | 8.8 | 25, 22, 46 | 399 |
| | ParaHippocampal_R | 8.8 | 22, -16, -22 | 63 |
| RSN 09 | Cingulum_Ant_L | 17 | -4, 49, 0 | 30768 |
| | Hippocampus_L | 9.9 | -23, -20, -16 | 127 |
| | Cingulum_Mid_L | 9.9 | -2, -40, 37 | 152 |
| | Cingulum_Mid_L | 9.4 | 0, -16, 37 | 104 |
| | Frontal_Sup_L | 9.2 | -22, 34, 46 | 387 |
| | Frontal_Inf_Orb_R | 9.2 | 36, 36, -12 | 123 |
| | Frontal_Inf_Orb_L | 9.1 | -32, 32, -14 | 119 |
| RSN 21 | Frontal_Inf_Orb_L | 11.6 | -50, 24, -2 | 28364 |
| | Supp_Motor_Area_L | 10.9 | -4, 10, 64 | 4018 |
| | Temporal_Mid_R | 8.6 | 48, -34, 0 | 1627 |
| | Precentral_L | 8.4 | -44, 4, 52 | 1137 |
| | Cerebellum_6_R | 8.2 | 22, -70, -28 | 531 |
| | Caudate_L | 7.6 | -15, 10, 10 | 401 |
| | Temporal_Pole_Mid_R | 7.2 | 50, 16, -24 | 632 |
| | Cerebellum_Crus2_R | 7.2 | 20, -78, -44 | 422 |
| | Frontal_Inf_Orb_R | 6.4 | 50, 30, -4 | 231 |
| RSN24 | Cingulum_Post_L | 20.4 | -4, -26, 30 | 8894 |

| | | | | |
|---|---|---|---|---|
| | Precuneus_R | 13.7 | 10, -67, 38 | 1971 |
| | Precuneus_L | 13.4 | -6, -68, 38 | 2049 |
| **Control System** | | | | |
| RSN 06 | Parietal_Inf_L | 10.8 | -54, -40, 46 | 4964 |
| | Frontal_Inf_Oper_L | 9.7 | -50, 8, 18 | 1386 |
| | Frontal_Inf_Tri_L | 9 | -44, 40, 16 | 3960 |
| | Temporal_Mid_L | 8.8 | -56, -60, -2 | 2272 |
| | Cingulum_Mid_L | 8.6 | -9, -30, 40 | 155 |
| | Parietal_Inf_R | 8.3 | 60, -34, 48 | 1076 |
| | Frontal_Mid_Orb_L | 8.2 | -28, 37, -12 | 1285 |
| | Parietal_Inf_L | 7.8 | -35, -46, 40 | 261 |
| | Frontal_Inf_Tri_R | 7.5 | 44, 40, 8 | 2242 |
| | Cerebellum_8_R | 6.4 | 22, -70, -50 | 156 |
| | Temporal_Mid_R | 6.3 | 60, -53, 2 | 238 |
| | Frontal_Inf_Orb_R | 6.3 | 24, 29, -14 | 109 |
| | Fusiform_L | 5.6 | -46, -54, -20 | 41 |
| RSN 07 | Angular_L | 9.8 | -40, -66, 52 | 10531 |
| | Cerebellum_Crus2_R | 8.6 | 42, -68, -42 | 4187 |
| | Frontal_Mid_L | 8.6 | -28, 20, 56 | 7511 |
| | Temporal_Mid_L | 8.3 | -64, -32, -8 | 6888 |
| | Frontal_Mid_Orb_L | 6.7 | -36, 54, -2 | 1739 |
| | Cerebellum_Crus1_R | 6.6 | 16, -84, -26 | 849 |
| RSN 08 | Parietal_Inf_R | 11.6 | 46, -54, 48 | 12370 |
| | Frontal_Mid_R | 9.7 | 42, 22, 46 | 12929 |
| | Temporal_Mid_R | 9.6 | 64, -30, -12 | 5797 |
| | Frontal_Mid_Orb_R | 9.3 | 40, 52, -2 | 7828 |
| | Frontal_Sup_Medial_R | 8.5 | 4, 30, 44 | 1274 |
| | Cerebellum_Crus2_L | 8.3 | -10, -80, -26 | 1251 |
| | Cerebellum_Crus1_L | 7.6 | -32, -66, -30 | 667 |
| | Cerebellum_Crus2_L | 7.3 | -38, -68, -45 | 1765 |
| RSN 20 | Frontal_Inf_Oper_L | 10.6 | -46, 12, 30 | 21076 |
| | Parietal_Inf_L | 9.8 | -36, -53, 44 | 4559 |
| | Cerebellum_Crus2_R | 9.1 | 10, -80, -32 | 2140 |
| | Frontal_Mid_L | 8.5 | -28, 12, 58 | 1116 |
| | Frontal_Sup_Medial_L | 8.3 | -4, 28, 44 | 1516 |
| | Temporal_Inf_L | 8.1 | -56, -50, -10 | 1680 |
| | Cerebellum_6_R | 7.6 | 28, -64, -30 | 678 |
| | Cerebellum_7b_R | 7.5 | 33, -70, -50 | 872 |

| | | | | |
|---|---|---|---|---|
| | Frontal_Inf_Tri_R | 6.7 | 48, 32, 22 | 1005 |
| | Cingulum_Ant_L | 6.5 | -4, 2, 28 | 106 |
| **Attention System** | | | | |
| RSN 11 | Precuneus_R | 12.1 | 6, -61, 58 | 10517 |
| | Frontal_Sup_R | 9.9 | 26, 4, 58 | 1855 |
| | Frontal_Sup_L | 9.2 | -24, 0, 62 | 1905 |
| | Occipital_Mid_R | 8.9 | 38, -81, 32 | 275 |
| | Cerebellum_9_R | 7.5 | 12, -50, -52 | 320 |
| | Cerebellum_9_L | 6.6 | -12, -50, -52 | 98 |
| RSN 12 | Parietal_Sup_R | 12.3 | 22, -64, 54 | 11630 |
| | Parietal_Sup_L | 11.4 | -21, -72, 48 | 9679 |
| | Precentral_R | 9.4 | 28, -4, 56 | 1102 |
| | Frontal_Sup_L | 8.6 | -26, -5, 58 | 575 |
| | Temporal_Inf_L | 8.4 | -52, -66, -6 | 47 |
| | Temporal_Inf_R | 7.6 | 56, -60, -11 | 135 |
| RSN 15 | Parietal_Sup_R | 16.2 | 22, -50, 66 | 6428 |
| | Parietal_Sup_L | 16.1 | -22, -50, 66 | 6758 |
| | Frontal_Sup_R | 11.6 | 20, -10, 72 | 1035 |
| | Frontal_Sup_L | 9.7 | -22, -8, 68 | 726 |
| RSN 18 | Parietal_Inf_L | 15.3 | -46, -32, 42 | 8917 |
| | SupraMarginal_R | 13.6 | 42, -32, 44 | 8356 |
| | Precentral_L | 11 | -52, 6, 30 | 1249 |
| | Precentral_R | 10.9 | 54, 8, 30 | 1362 |
| | Rolandic_Oper_L | 10.7 | -38, -5, 15 | 93 |
| | Precentral_R | 9.7 | 26, -10, 56 | 141 |
| | Temporal_Inf_R | 9.6 | 53, -60, -6 | 213 |
| | Temporal_Mid_L | 9.3 | -49, -64, 0 | 254 |
| | Cerebellum_8_R | 8.1 | 14, -72, -50 | 36 |
| **SAL System** | | | | |
| RSN 04 | Insula_R | 15 | 36, 4, 10 | 9086 |
| | Insula_L | 14 | -36, 2, 10 | 7427 |
| | Rolandic_Oper_R | 12.3 | 56, -24, 22 | 5460 |
| | Temporal_Sup_L | 11.5 | -54, -28, 20 | 3870 |
| | Cingulum_Mid_R | 11 | 6, 4, 42 | 2391 |
| | Cingulum_Mid_R | 9 | 14, -28, 40 | 378 |
| RSN 05 | Cingulum_Mid_R | 13.7 | 8, 12, 37 | 14941 |
| | Rolandic_Oper_R | 13.4 | 50, 8, 2 | 2889 |

| | | | | |
|---|---|---|---|---|
| | Anterior_Insula_L | 12.4 | -48, 8, 0 | 2252 |
| | Cerebellum_6_L | 11.3 | -30, -60, -24 | 1152 |
| | Precentral_R | 11.2 | 48, -4, 51 | 3609 |
| | Precentral_L | 9.5 | -48, -5, 50 | 2411 |
| | Pallidum_R | 9.2 | 22, 2, 6 | 1412 |
| | Putamen_L | 8.7 | -23, -2, 10 | 624 |
| | Thalamus_R | 8.5 | 12, -16, 8 | 174 |
| | Putamen_R | 8.3 | 30, -16, 2 | 59 |
| | Caudate_R | 8.1 | 16, -10, 20 | 282 |
| | Cerebellum_6_R | 7.8 | 30, -60, -24 | 170 |
| | Frontal_Mid_R | 7.7 | 34, 46, 30 | 237 |
| | Frontal_Mid_L | 7 | -34, 46, 28 | 48 |
| RSN 13 | Anterior_Insula_R | 14 | 33, 16, -10 | 6758 |
| | Cingulum_Ant_L | 13.5 | -9, 24, 26 | 16146 |
| | Anterior_Insula_L | 12.8 | -40, 15, -4 | 7197 |
| | Putamen_R | 8.3 | 18, 12, -4 | 282 |
| | Caudate_R | 8.1 | 12, 10, 6 | 324 |
| | Amygdala_L | 7.4 | -21, 4, -12 | 42 |
| **SM System** | | | | |
| RSN 03 | Temporal_Sup_L | 12.4 | -60, -12, 4 | 20386 |
| | Temporal_Sup_R | 11.9 | 66, -28, 6 | 21190 |
| RSN 16 | Paracentral_Lobule_L | 17.5 | -14, -28, 72 | 41543 |
| | Heschl_R | 12.3 | 36, -24, 18 | 672 |
| | Rolandic_Oper_L | 11.5 | -34, -26, 18 | 456 |
| | Putamen_L | 9.8 | -30, -14, 8 | 55 |
| | Putamen_R | 9.7 | 30, -11, 8 | 63 |
| | Cerebellum_4_5_R | 9.4 | 22, -36, -26 | 84 |
| RSN 17 | Postcentral_R | 19.6 | 46, -30, 58 | 18319 |
| | Cerebellum_6_L | 14.5 | -22, -53, -20 | 1471 |
| | Rolandic_Oper_R | 14 | 48, -22, 18 | 1176 |
| | Cingulum_Mid_R | 11.2 | 8, -24, 48 | 43 |
| | Cerebellum_8_L | 10.8 | -24, -54, -53 | 157 |
| RSN 19 | Postcentral_L | 18.8 | -40, -36, 63 | 18920 |
| | Cerebellum_6_R | 14.7 | 22, -50, -22 | 2420 |
| | Rolandic_Oper_L | 12.8 | -44, -22, 18 | 797 |
| | Supp_Motor_Area_L | 11.2 | -4, -12, 52 | 216 |
| | Cerebellum_8_R | 11 | 18, -58, -52 | 517 |
| | Vermis_6 | 10.5 | 5, -62, -16 | 202 |

| | | | | |
|---|---|---|---|---|
| | Cingulum_Mid_L | 10.3 | -6, -24, 46 | 39 |
| | Thalamus_L | 10.1 | -15, -22, 9 | 52 |
| RSN 22 | Postcentral_L | 19.8 | -50, -16, 34 | 15488 |
| | Postcentral_R | 19.2 | 52, -12, 32 | 14526 |
| | Cerebellum_6_L | 14.8 | -16, -62, -20 | 714 |
| | Cerebellum_6_R | 14.7 | 18, -60, -22 | 740 |
| | Pallidum_R | 13.6 | 28, -5, -8 | 463 |
| | Putamen_L | 13.1 | -29, -8, -6 | 499 |
| **VIS System** | | | | |
| RSN 10 | Temporal_Inf_R | 13.9 | 46, -70, -10 | 27515 |
| | Occipital_Inf_L | 13.6 | -44, -72, -8 | 25592 |
| RSN 14 | Calcarine_R | 19.4 | 16, -64, 10 | 68034 |
| | Vermis_7 | 11.3 | 2, -72, -26 | 272 |
| | Hippocampus_L | 10.5 | -20, -28, -4 | 46 |
| RSN 23 | Occipital_Inf_L | 15.8 | -32, -92, -8 | 9294 |
| | Occipital_Inf_R | 15.3 | 34, -88, -10 | 8339 |

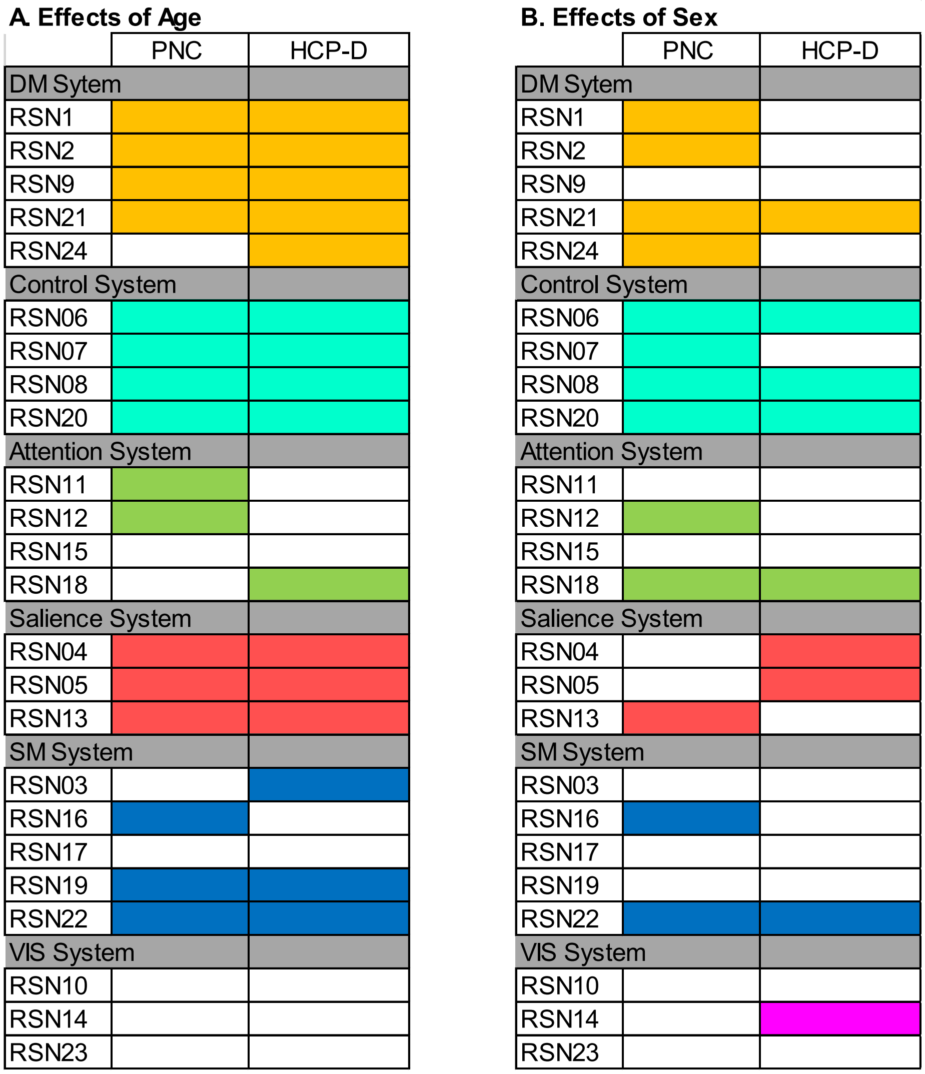

**Supplementary Figure S6: Summary of the effects of age and sex on spatial maps of each network, within the PNC and HCP-D Sample (for comparison), based on multivariate analyses**. Colored cases reflect networks with a significant effect at pFDR<0.01.

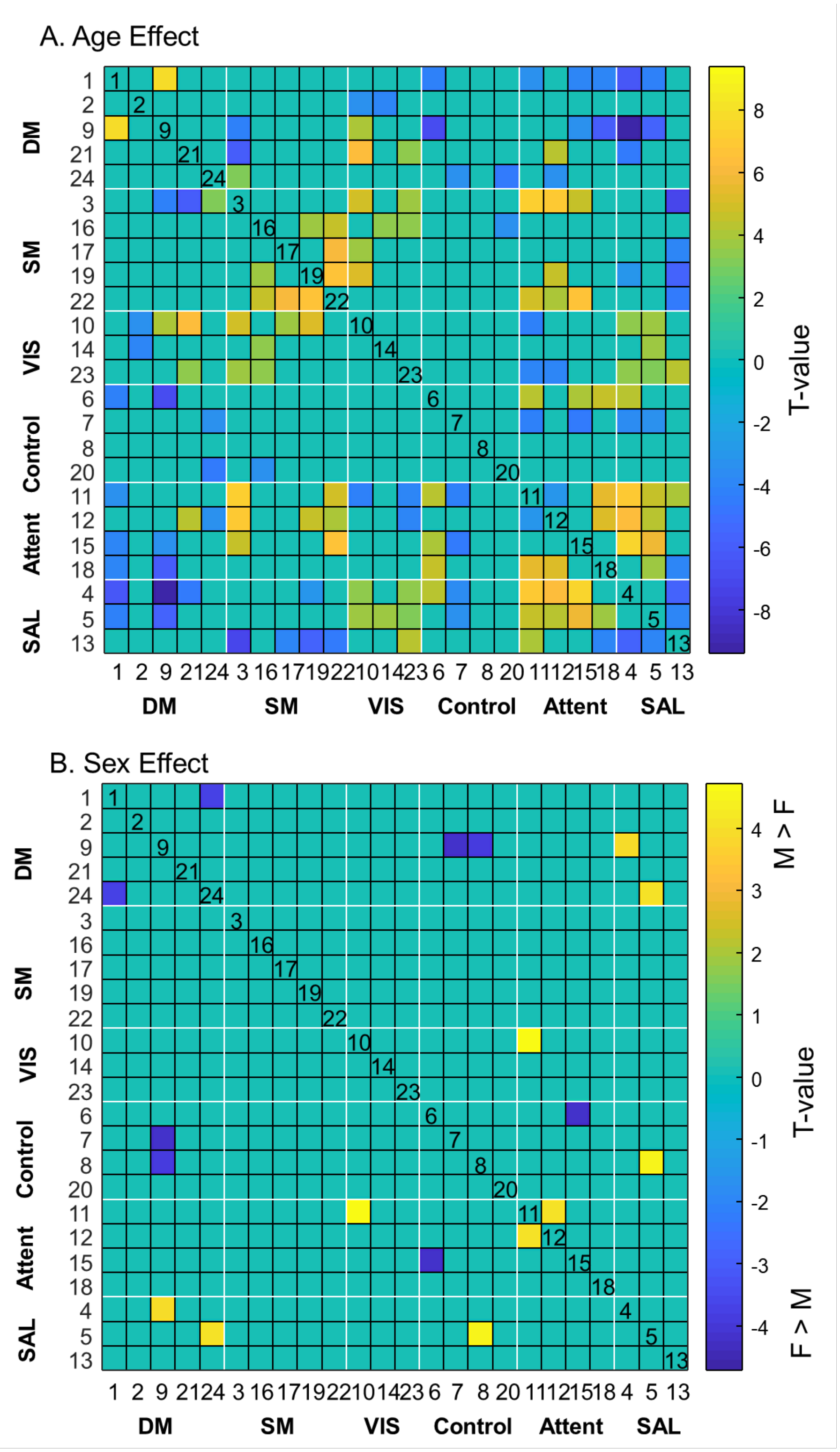

**Supplementary Figure S7: Effects of age and sex on FNC, within the HCP-D Sample**. Significant threshold was set at pFDR<0.01.

**Supplementary Table S3: Effects of Age on the spatial maps of Dev-Atlas RSNs.**

| | Brain Region | Effect direction | Peak | MNI coordinate (x,y,z) | Volume (mm3) |
|---|---|---|---|---|---|
| **DM System** | | | | | |
| RSN 01 | Frontal_Mid_Orb_R | positive | 15.7 | 3, 52, -1 | 16108 |
| | Frontal_Mid_R | positive | 5.9 | 28, 35, 46 | 2113 |
| | ParaHippocampal_R | positive | 5.6 | 17, -13, -22 | 653 |
| | ParaHippocampal_L | positive | 5.2 | -19, -21, -21 | 499 |
| | Lingual_L | positive | 4.3 | -15, -50, -2 | 454 |
| | Occipital_Mid_L | positive | 5.7 | -39, -83, 41 | 370 |
| | Frontal_Mid_L | positive | 4.3 | -26, 33, 46 | 355 |
| | Angular_L | negative | 11.3 | -45, -49, 26 | 10730 |
| | Angular_R | negative | 10.5 | 42, -49, 23 | 8269 |
| | Precuneus_R | negative | 9.7 | 8, -61, 27 | 4532 |
| | Temporal_Pole_Mid_L | negative | 9.7 | -39, 13, -35 | 1835 |
| | Olfactory_R | negative | 15.4 | 5, 9, -13 | 1273 |
| | Temporal_Mid_L | negative | 5.1 | -60, -9, -19 | 839 |
| | Temporal_Pole_Mid_R | negative | 6.2 | 33, 21, -36 | 775 |
| | Cerebelum_9_R | negative | 4.7 | 14, -47, -42 | 524 |
| | Cerebelum_Crus2_R | negative | 5.4 | 14, -83, -35 | 522 |
| RSN 02 | Precuneus_R | positive | 6.8 | 3, -58, 60 | 8690 |
| | Frontal_Mid_R | positive | 9 | 28, 32, 40 | 3748 |
| | Frontal_Mid_L | positive | 8.5 | -29, 35, 48 | 2579 |
| | Occipit_Mid_L | positive | 12.2 | -45, -81, 35 | 2347 |
| | Occipit_Mid_R | positive | 7.8 | 50, -75, 34 | 1676 |
| | Frontal_Mid_Orb_L | positive | 7.8 | -1, 54, -5 | 1571 |
| | Retrosplenial_R | negative | 12.1 | 19, -50, 13 | 12670 |
| | Fusiform_L | negative | 12.9 | -31, -41, -17 | 8738 |
| | Retrosplenial_L | negative | 11.4 | -15, -53, 12 | 4408 |
| | Occipital_Mid_R | negative | 6.6 | 33, -83, 23 | 2568 |
| | Olfactory_R | negative | 10.2 | 3, 12, -11 | 833 |
| | Temporal_Mid_R | negative | 4.8 | 53, -49, 20 | 565 |
| | Occipital_Mid_L | negative | 5.5 | -33, -85, 21 | 513 |
| | Occipital_Mid_L | negative | 4.6 | -39, -64, 26 | 418 |
| RSN 09 | Cingulum_Ant_L | positive | 13.1 | -3, 54, 4 | 2220 |
| | Frontal_Sup_L | positive | 5 | -17, 32, 55 | 509 |
| | Cingulum_Ant_R | positive | 4.1 | 3, 46, 13 | 242 |
| | Olfactory_L | negative | 15.7 | -1, 4, -13 | 5484 |
| | Frontal_Sup_Orb_L | negative | 6.8 | -17, 66, -2 | 863 |
| | Frontal_Sup_Med_R | negative | 6.6 | 2, 68, 1 | 488 |
| | ParaHippocampal_R | negative | 6.6 | 23, -22, -19 | 282 |

| | | | | | |
|---|---|---|---|---|---|
| | Rectus_R | negative | 5.5 | 8, 38, -19 | 280 |
| RSN 21 | Frontal_Sup_L | positive | 7.6 | -12, 46, 46 | 3879 |
| | Frontal_Inf_Orb_L | positive | 7.2 | -53, 38, -7 | 1714 |
| | Precentral_L | positive | 9 | -49, 7, 51 | 1343 |
| | Frontal_Sup_L | positive | 4 | -26, 49, 23 | 459 |
| | Frontal_Inf_Orb_R | positive | 5.6 | 53, 41, -7 | 400 |
| | Temporal_Mid_L | positive | 5.2 | -60, -64, 21 | 345 |
| | Supp_Motor_Area_R | positive | 5 | 9, 15, 68 | 304 |
| | Temporal_Pole_Mid_L | negative | 13 | -33, 19, -36 | 25171 |
| | Temporal_Pole_Mid_R | negative | 7.8 | 50, 12, -33 | 3596 |
| | Temporal_Sup_R | negative | 5.4 | 50, -30, 4 | 2403 |
| | Temporal_Sup_R | negative | 8.6 | 59, -11, -7 | 1424 |
| | Cerebelum_Crus1_R | negative | 5.2 | 11, -78, -25 | 712 |
| | Cerebelum_7b_R | negative | 5.8 | 16, -77, -45 | 660 |
| | Temporal_Inf_R | negative | 5.2 | 39, 4, -39 | 345 |
| RSN24 | n.s. | | | | |
| **Control System** | | | | | |
| RSN 06 | Frontal_Inf_Tri_L | positive | 9.2 | -46, 46, 9 | 3163 |
| | Frontal_Mid_R | positive | 7.9 | 50, 46, 4 | 1078 |
| | Frontal_Inf_Orb_L | positive | 7.8 | -31, 35, -13 | 902 |
| | Frontal_Mid_Orb_R | positive | 5.8 | 30, 41, -11 | 694 |
| | Frontal_Mid_Orb_L | positive | 4.8 | -46, 52, -2 | 393 |
| | Frontal_Mid_R | positive | 4.1 | 45, 43, 26 | 347 |
| | Frontal_Mid_L | positive | 5.2 | -33, 40, 43 | 333 |
| | Insula_R | positive | 4.4 | 39, 13, -5 | 290 |
| | Insula_L | positive | 5.6 | -39, 5, -1 | 226 |
| | Occipital_Mid_L | negative | 9.5 | -49, -71, -1 | 5395 |
| | SupraMarginal_L | negative | 8.6 | -54, -33, 32 | 2686 |
| | Temporal_Mid_R | negative | 5.6 | 53, -55, 6 | 1096 |
| | Frontal_Inf_Orb_L | negative | 8.8 | -21, 27, -19 | 693 |
| | Frontal_Inf_Orb_R | negative | 8.4 | 19, 24, -19 | 498 |
| | Cerebelum_8_R | negative | 6.6 | 17, -75, -53 | 431 |
| | SupraMarginal_R | negative | 5.8 | 47, -39, 34 | 337 |
| | SupraMarginal_R | negative | 6.2 | 56, -33, 48 | 296 |
| RSN 07 | Frontal_Mid_L | positive | 14.4 | -40, 60, 1 | 8775 |
| | Frontal_Sup_Medial_L | positive | 8.6 | -5, 35, 37 | 2233 |
| | Frontal_Mid_L | positive | 6.8 | -39, 15, 60 | 779 |
| | Frontal_Inf_Tri_L | positive | 4.7 | -46, 38, 27 | 604 |
| | Cerebelum_Crus2_R | negative | 10.6 | 30, -77, -45 | 10500 |

| | | | | | |
|---|---|---|---|---|---|
| | Angular_L | negative | 15.7 | -49, -53, 29 | 8773 |
| | Temporal_Inf_L | negative | 7.9 | -53, -27, -21 | 6624 |
| | Parietal_Inf_R | negative | 7.4 | 45, -55, 49 | 4916 |
| | Temporal_Mid_R | negative | 7.5 | 61, -29, -11 | 4218 |
| | Cerebelum_Crus2_L | negative | 7 | -37, -72, -45 | 1527 |
| | Precuneus_L | negative | 4.3 | -11, -64, 40 | 588 |
| | Cingulum_Mid_R | negative | 4.3 | 5, -44, 37 | 409 |
| | Precentral_L | negative | 4.4 | -33, 5, 49 | 238 |
| RSN 08 | Frontal_Mid_Orb_R | positive | 11.3 | 44, 55, -5 | 4237 |
| | Insula_R | positive | 8.2 | 36, 24, -5 | 1125 |
| | Cingulum_Ant_R | positive | 4.5 | 5, 43, 7 | 339 |
| | Frontal_Sup_Medial_R | positive | 5.6 | 3, 35, 57 | 277 |
| | Frontal_Sup_Medial_R | positive | 5 | 3, 33, 40 | 258 |
| | Cerebelum_Crus2_L | negative | 9.2 | -40, -72, -39 | 6621 |
| | Angular_L | negative | 5.7 | -46, -55, 37 | 2566 |
| | Angular_R | negative | 6.9 | 47, -57, 40 | 2346 |
| | Temporal_Inf_R | negative | 6.1 | 58, -41, -19 | 2321 |
| | Angular_R | negative | 5.1 | 39, -69, 40 | 1006 |
| | Frontal_Mid_Orb_R | negative | 5.6 | 28, 46, -2 | 715 |
| | Temporal_Mid_R | negative | 5.9 | 59, -44, -2 | 659 |
| | Precentral_R | negative | 5.2 | 36, 1, 49 | 392 |
| | Cerebelum_Crus2_L | negative | 3.8 | -5, -85, -31 | 202 |
| RSN 20 | Frontal_Mid_Orb_L | positive | 12.1 | -45, 52, -1 | 4202 |
| | Cerebelum_Crus2_R | negative | 8.5 | 8, -83, -29 | 774 |
| | Temporal_Inf_L | negative | 3.8 | -57, -53, -13 | 216 |
| **Attention System** | | | | | |
| RSN 11 | Lingual_L | positive | 6.7 | -17, -57, 4 | 2193 |
| | Lingual_R | positive | 5.6 | 8, -49, 6 | 587 |
| | Occipital_Mid_R | positive | 6.1 | 44, -81, 32 | 466 |
| | Occipital_Mid_L | positive | 6.5 | -40, -85, 34 | 404 |
| | Frontal_Mid_L | positive | 5.2 | -33, 41, 40 | 330 |
| | Frontal_Sup_L | positive | 6 | -26, 12, 68 | 313 |
| | Cuneus_R | positive | 4.7 | 19, -64, 26 | 300 |
| | Precuneus_R | negative | 5.9 | 9, -49, 55 | 223 |
| RSN 12 | Fusiform_R | positive | 5.6 | 23, -63, -11 | 1010 |
| | Frontal_Mid_R | positive | 6.6 | 33, -1, 63 | 438 |
| | Fusiform_L | positive | 5.2 | -31, -64, -15 | 330 |
| | Fusiform_L | positive | 4.8 | -25, -55, -13 | 310 |
| | Occipital_Sup_L | negative | 8.9 | -25, -63, 27 | 2639 |

| | | | | | |
|---|---|---|---|---|---|
| | Precuneus_R | negative | 11.7 | 19, -63, 41 | 2076 |
| | Occipital_Sup_R | negative | 5.2 | 25, -72, 29 | 415 |
| RSN 15 | n.s. | | | | |
| RSN 18 | n.s. | | | | |
| **SAL System** | | | | | |
| RSN 04 | Postcentral_L | positive | 6.8 | -25, -43, 71 | 1515 |
| | Insula_R | positive | 7 | 45, 21, -5 | 765 |
| | Frontal_Mid_R | positive | 8 | 33, 47, 27 | 711 |
| | Parietal_Inf_L | positive | 5.6 | -59, -30, 48 | 346 |
| | Rolandic_Oper_R | negative | 13.4 | 45, -25, 23 | 2555 |
| | Postcentral_L | negative | 8.1 | -57, -15, 20 | 1433 |
| | Insula_R | negative | 9.1 | 33, -16, 9 | 1054 |
| | Rolandic_Oper_R | negative | 5.7 | 47, -2, 12 | 288 |
| RSN 05 | Frontal_Mid_L | positive | 5.2 | -29, 49, 37 | 537 |
| | SupraMarginal_R | positive | 6.2 | 47, -33, 26 | 423 |
| | Putamen_L | positive | 5.8 | -26, -1, 4 | 366 |
| | Frontal_Mid_R | positive | 5.4 | 36, 52, 27 | 259 |
| | Putamen_R | positive | 7.6 | 28, -1, -7 | 228 |
| | Precentral_R | negative | 6.7 | 36, -15, 54 | 366 |
| | Sup_Motor_Area_R | negative | 5.8 | 16, -9, 55 | 259 |
| RSN 13 | Cingulum_Ant_L | positive | 10.9 | -3, 47, 1 | 4527 |
| | Frontal_Mid_R | positive | 7.8 | 28, 57, 26 | 1167 |
| | Thalamus_L | positive | 6.3 | -3, -5, 4 | 1055 |
| | Frontal_Mid_L | positive | 5.9 | -29, 57, 20 | 902 |
| | Supp_Motor_Area_R | positive | 7.5 | 3, 18, 68 | 466 |
| | Cingulum_Mid_R | positive | 5.1 | 3, -19, 46 | 445 |
| | Cingulum_Ant_L | positive | 4.8 | -3, 33, 20 | 425 |
| | Putamen_L | positive | 6.8 | -17, 7, -5 | 360 |
| | Frontal_Mid_L | positive | 5.4 | -29, 43, 37 | 334 |
| | Cingulum_Mid_L | positive | 5 | -5, -39, 43 | 311 |
| | Olfactory_R | negative | 10.1 | 14, 13, -13 | 3009 |
| | Olfactory_L | negative | 7.7 | -15, 9, -15 | 1507 |
| | Cingulum_Mid_L | negative | 6.6 | -15, 13, 32 | 825 |
| | Putamen_L | negative | 6 | -26, 18, 6 | 799 |
| | Cingulum_Ant_L | negative | 7.8 | 5, 15, 21 | 729 |
| | Frontal_Inf_Oper_R | negative | 5.1 | 39, 15, 12 | 680 |
| | Cingulum_Mid_R | negative | 5.7 | 11, 24, 37 | 642 |
| | Cingulum_Ant_L | negative | 5.9 | -6, 18, 20 | 307 |

| | | | | | |
|---|---|---|---|---|---|
| | Insula_L | negative | 4.7 | -35, 12, -7 | 226 |
| | Caudate_R | negative | 4.8 | 3, 7, -7 | 224 |
| **SM System** | | | | | |
| RSN 03 | n.s. | | | | |
| RSN 16 | Precentral_R | positive | 6.8 | 44, -13, 60 | 2317 |
| | Postcentral_R | positive | 5.4 | 39, -36, 63 | 397 |
| | Temporal_Sup_R | positive | 4.7 | 51, -35, 23 | 225 |
| | Precentral_L | positive | 6.7 | -33, -19, 71 | 216 |
| | Paracentral_Lobule_L | negative | 9.6 | -15, -35, 71 | 3193 |
| | Postcentral_R | negative | 8.8 | 16, -35, 69 | 820 |
| RSN 17 | n.s. | | | | |
| RSN 19 | Precentral_R | positive | 8.6 | 37, -15, 49 | 5135 |
| | Supp_Motor_Area_R | positive | 5.1 | 5, -16, 49 | 568 |
| | Cerebelum_8_R | negative | 7 | 16, -58, -50 | 1155 |
| | Putamen_L | negative | 7.4 | -33, -11, -2 | 1141 |
| | Postcentral_L | negative | 7.4 | -43, -30, 57 | 1062 |
| | Thalamus_L | negative | 6.4 | -11, -25, 9 | 406 |
| RSN 22 | Temporal_Sup_L | positive | 4.8 | -59, -39, 18 | 997 |
| | Paracentral_L | positive | 4.4 | -17, -29, 60 | 436 |
| | Supp_Motor_Area_L | positive | 4.1 | -6, -13, 62 | 371 |
| | Precentral_R | positive | 4.5 | 42, -9, 57 | 249 |
| | Postcentral_R | negative | 11 | 42, -7, 29 | 2546 |
| | Precentral_L | negative | 9.8 | -49, -5, 26 | 1372 |
| | Pallidum_R | negative | 8.6 | 30, -13, -7 | 406 |
| | Hippocampus_L | negative | 6.1 | -29, -9, -11 | 370 |
| | Thalamus_L | negative | 7.8 | -12, -22, 1 | 313 |
| **VIS System** | | | | | |
| RSN 10 | n.s. | | | | |
| RSN 14 | n.s. | | | | |
| RSN 23 | n.s. | | | | |

Only clusters > 200mm$^3$ are listed. Clusters information was extracted through MRIcroGL.

**Supplementary Table S4: Effects of sex on the spatial maps of Dev-Atlas RSNs.**

| | Brain Region | Effect direction | Peak | MNI coordinate (x,y,z) | Volume (mm3) |
|---|---|---|---|---|---|
| **DM System** | | | | | |
| RSN 01 | Temporal_Mid_L | M>F | 9.5 | -59, -61, 23 | 5315 |
| | Temporal_Mid_R | M>F | 9.5 | 56, -58, 13 | 3018 |
| | Temporal_Inf_R | M>F | 4.2 | 58, -9, -31 | 263 |
| | Frontal_Inf_Orb_L | M>F | 6.5 | -33, 32, -13 | 217 |
| | Rectus_R | F>M | 8.4 | 2, 33, -21 | 2758 |
| | Precuneus_R | F>M | 7.1 | 5, -57, 23 | 546 |
| | Temporal_Mid_R | F>M | 5.5 | 47, -16, -11 | 510 |
| | Frontal_Sup_Medial_L | F>M | 5.8 | -11, 68, 1 | 352 |
| RSN 02 | Temporal_Mid_R | M>F | 7 | 47, -67, 18 | 3873 |
| | Occipital_Mid_L | M>F | 7.4 | -45, -78, 29 | 3503 |
| | Precuneus_L | M>F | 4.8 | -6, -63, 57 | 677 |
| | Cingulum_Mid_L | M>F | 5.6 | -6, -35, 43 | 632 |
| | ParaHippocampal_R | F>M | 9.6 | 19, -19, -17 | 3164 |
| | Temporal_Mid_R | F>M | 6.2 | 47, -15, -13 | 880 |
| | Precuneus_R | F>M | 4.7 | 22, -58, 27 | 868 |
| | ParaHippocampal_L | F>M | 5.1 | -19, -22, -15 | 866 |
| | Frontal_Mid_R | F>M | 7.6 | 28, 33, 32 | 721 |
| | Lingual_R | F>M | 5.6 | 9, -49, 4 | 686 |
| | Fusiform_L | F>M | 4.5 | -17, -39, -11 | 569 |
| | Calcarine_L | F>M | 5.3 | -11, -49, 6 | 348 |
| | Precuneus_L | F>M | 6.6 | -11, -47, 51 | 260 |
| RSN 09 | n.s. | | | | |
| RSN 21 | Frontal_Sup_L | F>M | 8.9 | -17, 49, 37 | 5491 |
| | Temporal_Inf_R | F>M | 7.3 | 47, 4, -33 | 846 |
| | Frontal_Inf_Orb_R | F>M | 6.3 | 45, 40, -5 | 571 |
| | Temporal_Mid_R | F>M | 5.6 | 44, -43, 7 | 544 |
| | Temporal_Mid_L | F>M | 6.1 | -45, -39, 1 | 482 |
| | Supp_Motor_Area_R | F>M | 4.5 | 14, 18, 65 | 406 |
| | Frontal_Inf_Oper_L | F>M | 4.9 | -53, 13, 20 | 339 |
| | Temporal_Inf_L | F>M | 5 | -45, 9, -41 | 284 |
| RSN24 | Cuneus_L | M>F | 11.6 | -3, -75, 32 | 3617 |
| | Angular_L | M>F | 3.9 | -49, -61, 37 | 231 |
| | Cingulum_Post_R | F>M | 12.7 | 3, -39, 27 | 8577 |
| | Precuneus_L | F>M | 4.9 | -12, -50, 35 | 750 |
| | Cingulum_Post_R | F>M | 5.2 | 2, -39, 9 | 328 |

| | | | | | |
|---|---|---|---|---|---|
| **Control System** | | | | | |
| RSN 06 | Frontal_Mid_R | F>M | 10.8 | 37, 41, 12 | 1265 |
| | Frontal_Inf_Orb_L | F>M | 7 | -29, 33, -13 | 528 |
| | Frontal_Inf_Orb_R | F>M | 5.5 | 25, 32, -11 | 250 |
| RSN 07 | SupraMarginal_L | M>F | 8.3 | -63, -49, 32 | 5942 |
| | Cerebellum_Crus2_R | M>F | 8.8 | 17, -86, -29 | 1048 |
| | Precuneus_R | M>F | 5 | 2, -57, 41 | 376 |
| | Angular_R | M>F | 5 | 42, -57, 27 | 234 |
| | Frontal_Mid_L | F>M | 6.5 | -25, 21, 46 | 1437 |
| | Frontal_Sup_L | F>M | 5.2 | -21, 61, 4 | 447 |
| | Temporal_Mid_L | F>M | 5.1 | -60, -43, 6 | 259 |
| | Frontal_Sup_Orb_L | F>M | 4.7 | -25, 47, -11 | 209 |
| RSN 08 | Angular_R | M>F | 8.8 | 51, -63, 48 | 9495 |
| | Cerebellum_Crus1_L | M>F | 5.4 | -19, -85, -29 | 1118 |
| | Cerebellum_7b_L | M>F | 8.3 | -45, -64, -55 | 442 |
| | Frontal_Mid_R | M>F | 4.2 | 47, 41, 23 | 357 |
| | Frontal_Sup_R | M>F | 5.4 | 22, 55, 35 | 330 |
| | Temporal_Inf_R | M>F | 4.1 | 64, -53, -8 | 283 |
| | Frontal_Mid_R | M>F | 5.2 | 42, 54, 18 | 259 |
| | Frontal_Mid_Orb_R | F>M | 6 | 42, 52, -2 | 723 |
| | Frontal_Sup_Medial_L | F>M | 4.4 | -1, 27, 51 | 377 |
| RSN 20 | Cerebellum_Crus1_R | M>F | 5.8 | 16, -81, -29 | 251 |
| | Frontal_Inf_Oper_L | F>M | 5.6 | -35, 9, 29 | 706 |
| **Attention System** | | | | | |
| RSN 11 | n.s. | | | | |
| RSN 12 | Angular_L | F>M | 6.7 | -31, -53, 37 | 563 |
| RSN 15 | n.s. | | | | |
| RSN 18 | Rolandic_Oper_L | M>F | 7.8 | -57, 4, 13 | 312 |
| | Postcentral_R | M>F | 5.5 | 67, -15, 21 | 212 |
| | SupraMarginal_L | F>M | 8.4 | -46, -25, 35 | 913 |
| | Parietal_Inf_L | F>M | 6.9 | -37, -36, 32 | 342 |
| | Frontal_Mid_R | F>M | 6.1 | 44, 35, 20 | 302 |
| **SAL System** | | | | | |
| RSN 04 | n.s. | | | | |

| | | | | | |
|---|---|---|---|---|---|
| RSN 05 | n.s. | | | | |
| RSN 13 | Temporal_Pole_Sup_R | M>F | 7.5 | 50, 18, -11 | 1800 |
| | Temporal_Pole_Sup_L | M>F | 7.1 | -49, 18, -15 | 1468 |
| | Insula_R | M>F | 6.9 | 31, 13, -13 | 230 |
| | Caudate_L | F>M | 6.1 | -12, 7, 9 | 1345 |
| | Thalamus_L | F>M | 7.4 | -5, -15, 13 | 1273 |
| | Caudate_R | F>M | 5.3 | 9, 7, -11 | 691 |
| | Cingulum_Ant_L | F>M | 5.1 | -9, 24, 32 | 211 |
| **SM System** | | | | | |
| RSN 03 | n.s. | | | | |
| RSN 16 | Paracentral_Lobule_R | M>F | 8 | 8, -41, 71 | 617 |
| RSN 17 | n.s. | | | | |
| RSN 19 | n.s. | | | | |
| RSN 22 | Insula_L | F>M | 9.1 | -40, -11, 15 | 997 |
| | Postcentral_R | F>M | 7.1 | 50, -2, 27 | 440 |
| **VIS System** | | | | | |
| RSN 10 | n.s. | | | | |
| RSN 14 | n.s. | | | | |
| RSN 23 | n.s. | | | | |

Only clusters > 200mm$^3$ are listed. Clusters information was extracted through MRIcroGL.
